# Allele-specific NKX2-5 binding underlies multiple genetic associations with human EKG traits

**DOI:** 10.1101/351411

**Authors:** Paola Benaglio, Agnieszka D’Antonio-Chronowska, William W. Greenwald, Christopher DeBoever, He Li, Frauke Drees, Sanghamitra Singhal, Hiroko Matsui, Matteo D’Antonio, Erin N. Smith, Kelly A. Frazer

## Abstract

Genetic variation affecting the binding of transcription factors (TFs) has been proposed as a major mechanism underlying susceptibility to common disease. NKX2-5, a key cardiac development TF, has been associated with electrocardiographic (EKG) traits through GWAS, but the extent to which differential binding of NKX2-5 contributes to these traits has not yet been studied. Here, we analyzed transcriptomic and epigenomic data generated from iPSC-derived cardiomyocyte lines (iPSC-CMs) from seven whole-genome sequenced individuals in a three-generational family. We identified ~2,000 single nucleotide variants (SNVs) associated with allele-specific effects (ASE) on NKX2-5 binding. These ASE-SNVs were enriched for altered TF motifs (both cognate and other cardiac TFs), and were positively correlated with changes in H3K27ac in iPSC-CMs, suggesting they impact cardiac enhancer activity. We found that NKX2-ASE-SNVs were significantly enriched for being heart-specific eQTLs and EKG GWAS variants, suggesting that altered NKX2-5 binding at multiple sites across the genome influences EKG traits. We used a fine-mapping approach to integrate iPSC-CM molecular phenotype data with a GWAS for heart rate, and determined that NKX2-5 ASE variants are likely causal for numerous known, as well as previously unidentified, heart rate loci. Analyzing Hi-C and gene expression data from iPSC-CMs at these heart rate loci, we identified several genes likely to be causally involved in heart rate variability. Our study demonstrates that differential binding of NKX2-5 is a common mechanism underlying genetic association with EKG traits, and shows that characterizing variants associated with differential binding of development TFs in iPSC-derived cell lines can identify novel loci and mechanisms influencing complex traits.

## INTRODUCTION

Altered transcription factor (TF) binding has been proposed as one of the major mechanisms by which non-coding regulatory variants are causally associated with complex traits^1–4^. Additionally, it has been hypothesized that all genetic variation that is functional in a disease-associated cell type may be relevant for the disease^5^. These two postulates suggest that characterizing regulatory variation that alters TF binding in cardiomyocytes could not only reveal the molecular effects of variants identified to date through cardiac trait genome-wide association studies (GWAS), but also enable the identification of novel functional variants underlying these cardiac phenotypes. Recent GWAS of electrocardiographic (EKG) phenotypes^6–11^ have found >500 risk variants, the majority of which are non-coding. Detecting the causal variants that drive these associations, and systematically interrogating their molecular mechanisms, is challenging because functional regulatory variation: 1) is frequently associated with many other neutral variants due to linkage disequilibrium (LD)^12^; 2) can regulate genes that are distant from their location; and 3) often functions in a cell type-specific manner^13–15^. Furthermore, despite the fact that the most recent GWAS for EKG traits have typically involved more than 20,000 individuals, only a fraction of the heritability is explained by the identified risk variants^16,17^. Thus, there are likely hundreds of additional variants that contribute to variability in EKG traits across individuals that will be difficult to detect through GWAS. For these reasons, experimental approaches that enable genome-wide characterization of regulatory variation in cardiomyocytes are needed.

Regulatory genetic studies are currently being performed by large consortia^18^ using adult heart tissues; however, it is typically difficult to get post-mortem samples, the amount of material obtained is limited and hence multiple molecular assays often cannot be conducted on the same sample, and the epigenetic states of the tissue may be affected by lifetime exposures. On the other hand, human induced pluripotent stem cell (iPSC) derived cardiomyocytes (iPSC-CMs) could be an advantageous model system because samples with different genetic backgrounds are readily obtained, iPSCs provide a renewable source of derived cell types and hence many molecular assays can be performed on the same sample, and the iPSC reprogramming process eliminates most epigenetic memory^19^. Genetic variation has been shown to affect gene expression and DNA methylation in undifferentiated iPSCs, with most variation between cell lines being explained by their genetic background^19–21^. Additionally, multiple studies have shown the utility of iPSC collections and derived cell types to perform expression QTL (eQTL) studies^20–25^. However, there are only a few studies showing similar utility of iPSC-CMs to study regulatory variants relevant for molecular phenotypes^23^. Despite the many advantages of using iPSC-CMs for genetic studies, there are concerns about the cell type composition variability between cell lines that arises from directed differentiation^25,26^. Another concern is that, because cultured cells are imperfect models of primary tissues, not all genetic variants that alter cardiac phenotypes in adult heart tissues will be detected in iPSC-CMs, which tend to have molecular and functional proprieties similar to fetal heart cells^27^. Thus, while human iPSC-CMs are a promising model system, it has yet to be shown that they could enable the identification and characterization of regulatory variants that play important roles in cardiac traits and disease.

Mammalian heart development, which has been primarily studied in the mouse, is regulated by evolutionary conserved transcription factors (TFs) including NKX2-5, TBX5, GATA4, MEF2A, SRF and MEIS1. In the murine model system, these TFs have been shown to cooperatively bind to common enhancer and promoter regions^28–32^. Coding mutations in *NKX2-5, TBX5*, and *GATA4* cause a spectrum of human congenital heart defects33, and common non-coding DNA variants near *NKX2-5, TBX5*, and *MEIS1* genes have been associated with EKG phenotypes through GWAS6,10,34. Furthermore, the most recent atrial fibrillation GWAS identified associated loci that are important in development of the fetal heart, indicating that variation in developmental pathways may play an important role in the etiology of EKG phenotypes35. Therefore, it is likely that genetic variation affecting the binding of developmental cardiac TFs also influences the heritability of EKG traits. However, this hypothesis has not yet been demonstrated on a genome-wide scale, and the extent to which differential TF binding underlies these cardiac phenotypes is unknown.

Here, we conducted a genome-wide analysis to identify regulatory variants affecting the binding of NKX2-5, and investigated their role in cardiac gene expression and EKG phenotypes. We generated iPSC-CM lines from a pedigree of seven whole-genome sequenced individuals, and profiled them with a variety of functional genomic assays including RNA-Seq, ATAC-Seq, and ChIP-Seq of both histone modification H3K27ac and NKX2-5. First, to establish the robustness of the cellular model, we evaluated how well iPSC-CMs recapitulate cardiomyocyte-specific expression and epigenetic signatures, and how genetic variants effected the variability of these molecular phenotypes across the iPSC-CM lines. Then, for each individual, we identified heterozygous sites that showed allele-specific effects (ASE) in the molecular phenotypes, and investigated NKX2-5 ASE variants in detail by examining whether they altered cardiac TF motifs, or were eQTLs or EKG GWAS-SNPs. To evaluate the extent to which NKX2-5 ASE could be used to prioritize causal EKG variants, we then integrated iPSC-CM molecular phenotype data with summary statistics from a GWAS meta-analysis for heart rate using a fine-mapping approach. We found that genetic variation affecting the binding of NKX2-5 and other cardiac TFs (with which NKX2-5 cooperatively binds) likely underlies the molecular mechanisms of numerous heart rate loci across the genome, including previously unidentified loci. Finally, by analyzing Hi-C chromatin interaction data from these iPSC-CMs, and gene expression data from 128 iPSC-CMs, we identified potential target genes for several putative causal NKX2-5 ASE variants influencing heart rate at both known and novel genetic loci.

## RESULTS

### Generation of iPSC-derived cardiomyocytes

We generated iPSC-derived cardiomyocytes (iPSC-CMs) from seven individuals in a three-generation family that includes three genetically unrelated subjects and two parent-offspring quartets (**Fig. 1a; Supplemental Table 1**). Fibroblasts from these individuals were reprogrammed using Sendai virus, and nine iPSC lines previously shown to be pluripotent and to have high genomic integrity^36^ were obtained. Using a monolayer cardiac differentiation approach^37^, we differentiated the nine iPSCs into 26 iPSC-CM lines: 12 were harvested at day 25 after lactate selection to obtain purer cardiomyocytes, and 14 were harvested at day 15, of which one was lactate purified (**Fig. 1a**). All iPSC-CMs displayed synchronous, spontaneous beating, and, on average, 89% +/− 8.4% of the cells in the lactate purified iPSC-CM lines and 68.6% +/− 13% in the non-lactate purified iPSC-CM lines stained positive for the cardiomyocyte-specific marker cardiac troponin T (TNNT2) by flow cytometry (**Supplementary Table 2, Supplementary Fig 1a**). Immunofluorescence for TNNT2 and MYL7 on iPSC-CM lines confirmed that they developed cardiomyocyte-like cytoskeletal structures (**Supplementary Fig 1b**). These data show that the derived iPSC-CMs display contractile activity and morphology similar to human cardiomyocytes.

**Figure 1.**
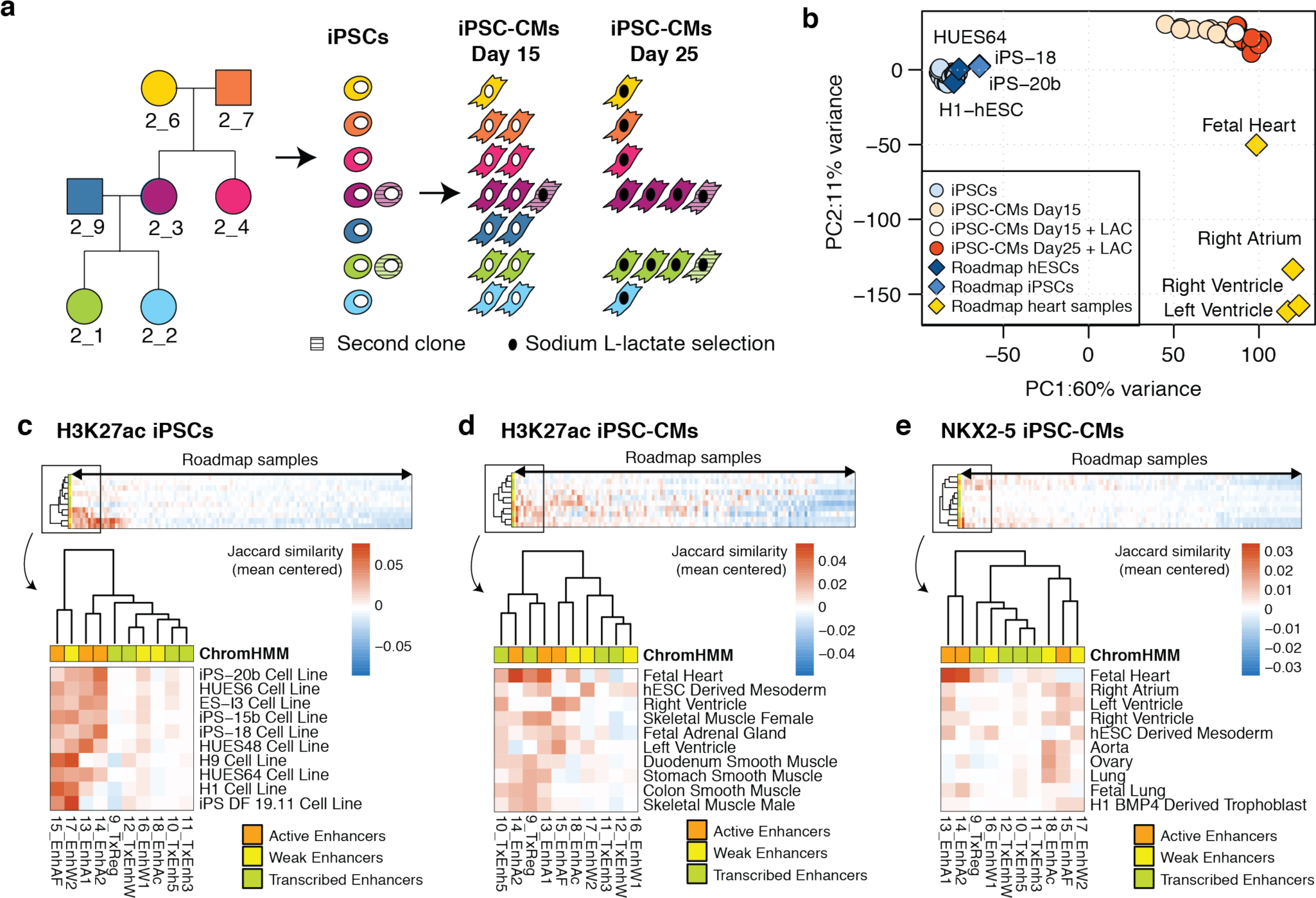
Generation and characterization of iPSCs and iPSC-CMs by gene expression and epigenetic profiling. (**a**) Pedigree showing the relationships of the seven individuals. Biological replicates of iPSC-CM lines were derived from independent differentiations of one or two iPSC clones from each individual (the second clone from 2_3 and 2_1 is indicated by hatches). iPSC-CMs harvested at day 25 and one of those harvested at day 15 underwent a cardiomyocyte purification step using L-lactate media (indicated by black filled circles). (**b**) Principal component 1 and 2 of RNA-Seq (15,725 genes) from iPSCs (29 samples from 7 individuals), iPSC-CMs (27 samples from 7 individuals), Roadmap stem cell lines (H1, HUES64, iPS-20b and iPS18) and human tissues (right ventricle, left ventricle, right atrium and fetal heart). (**c-e**) Heatmap and hierarchical clustering of similarity based on overlap between enhancer annotations (from 25-state ChromHMM) for 127 samples from Roadmap Epigenomics^14,40^ and ChIP-Seq peaks: (**c**) H3K27ac in iPSCs, (**d**) iPSC-CMs and (**e**) NKX2-5 in iPSC-CMs. Similarity was calculated using the *Jaccard* statistic (intersection/union of base pairs in each comparison), mean-centered across the 127 tissues and ordered by highest average enhancer similarity. A zoom in of the top 10 Roadmap tissues with highest average enhancer similarity is shown.

### Functional genomic profiles of iPSC-CMs recapitulate cardiomyocyte-specific characteristics

We investigated if the genome-wide gene expression and regulatory activity of the iPSC-CM lines were more similar to cardiac tissues than other human tissues, or the iPSCs from which they were derived. We generated and analyzed 56 RNA-Seq (iPSCs: 29 biological replicates; iPSC-CMs: 26 biological and 1 technical replicates), 21 ATAC-Seq (iPSCs: 10 biological replicates; iPSC-CMs: 10 biological and 1 technical replicates), 48 ChIP-Seq of histone modification H3K27ac (iPSCs: 17 biological and 4 technical replicates; iPSC-CMs: 25 biological and 2 technical replicates), and 15 ChIP-seq of NKX2-5 (iPSC-CMs: 12 biological and 3 technical replicates) (**Supplementary Table. 2** and 3, **Supplementary Fig. 2**). iPSC-CMs and iPSCs, respectively, expressed cardiac-specific and stem cell genes^38,39^ (**Supplementary Fig. 1c**). Additionally, we observed coordinated changes in TF binding and chromatin activity at the transcription start site of genes differentially expressed between these two cell types (**Supplementary Fig. 3a**). Principal component (PC) analysis of genome-wide gene expression further showed that iPSC-CMs and iPSCs clustered separately from each other, and closer to human cardiac tissue and stem cell reference samples^14^, respectively (**Fig. 1b**), with the iPSC-CMs that were lactate purified having similar PC1 values to fetal heart tissue. To identify which tissues among those in the 127 Roadmap Epigenomics^14,40^ were the most similar to the iPSC-CMs and iPSCs, we examined the enhancers of each Roadmap tissue, as enhancers are known to be highly tissue-specific. Specifically, we determined the overlap between the ten enhancer annotations from 25-state ChromHMM, and either ChIP-Seq peaks (**Fig. 1c-e**) or ATAC-Seq peaks (**Supplementary Fig. 3b-c**). As expected, the tissues with the highest degree of enhancer similarity with ChIP-Seq and ATAC-Seq were fetal heart for iPSC-CMs, and stem cells for iPSCs. Notably, cardiac and mesodermal tissues were also among the top 10 tissues most similar to iPSC-CMs. These analyses show that the genomic datasets generated for the iPSC-CMs recapitulate cardiomyocyte-specific gene expression and regulatory elements.

### Genetic background underlies variability of molecular phenotypes in iPSC-CMs

Experimental sources of variation across the iPSC-CMs, such as differentiation efficiency, may confound the effects that are driven by different genetic backgrounds, thereby limiting our ability to use these cell lines as a genetic model^25^. To identify the sources of variability in our iPSC-CM datasets, and evaluate the contribution of the genetic background to this variation compared with the iPSCs, we performed PC analysis on each of the RNA-Seq and ChIP-Seq datasets, and tested whether known covariates, such as batch, TNNT2 expression (for iPSC-CMs), and subject, were associated with each of the top 10 PCs. While we observed variation in both the iPSC-CMs and iPSCs due to differentiation efficiencies and/or batch effects (**Supplementary Fig. 4**), the average sample-to-sample Spearman correlation of molecular phenotypes was higher between samples of the same individual, than between different individuals (technical replicates were not compared with each other) (Mann Whitney test *P*< 0.05); additionally, samples of related individuals tended to be more correlated than samples of unrelated individuals (**Fig. 2**). Of note, the iPSC-CMs showed slightly greater variation (i.e. lower correlation values) than the iPSCs, likely due to cellular heterogeneity as previously observed^25,26^. Overall, ^1^ < these analyses indicate that genetic background was a major driver of variability in the iPSC-CM molecular datasets, and further support the use of the iPSC-CMs as a model system for interrogating the functions of regulatory variants.

**Figure 2.**
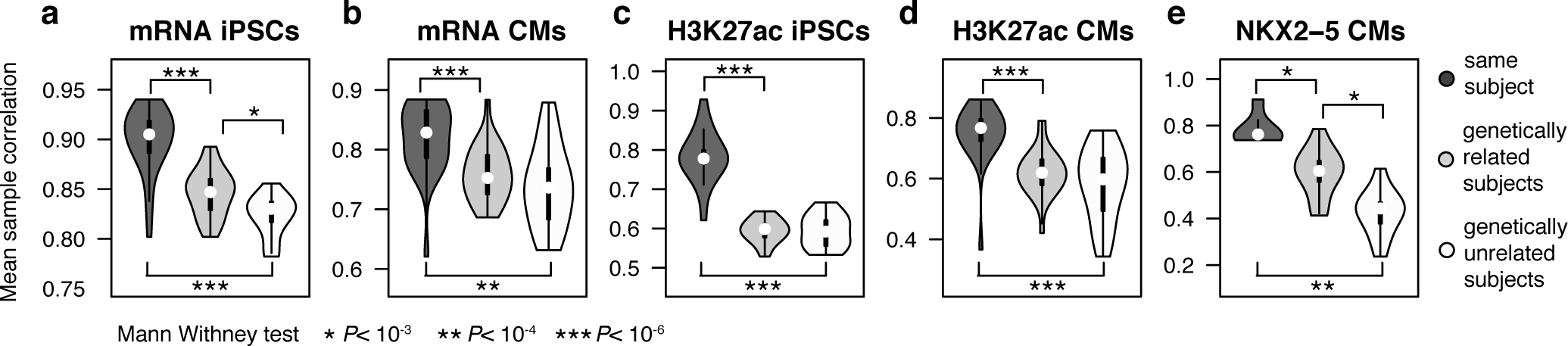
Correlation of molecular phenotypes between samples with different genetic relationships. Distributions of the average Spearman correlation coefficients between pairs of samples across the 1,000 most variable genes (**a**,**b**) or peaks (**c-e**) for the indicated molecular phenotypes. The average per-sample pairwise correlation was calculated between samples from either the same subject, genetically related subjects or genetically unrelated subjects. Technical replicates were excluded for pair-wise comparisons between samples of the same subject.

### NKX2-5 peaks commonly show allele-specific effects

We examined the fraction of iPSC-CM NKX2-5 peaks affected by genetic variants compared with the fraction of affected H3K27ac peaks and expressed genes (in both iPSC-CMs and iPSCs) by identifying heterozygous sites that showed allele-specific effects (ASE) in each individual. When multiple individuals carried the same heterozygous SNV, we combined the ASE results across the samples in a meta-analysis. For each phenotype, we tested between 19,371 (NKX2-5) to 123,151 (H3K27ac in iPSC-CMs) heterozygous SNVs within 12,492 to 57,631 regions (genes or peaks) (**Fig. 3a**), and identified between 375 (H3K27ac in iPSCs) and 1,941 (NKX2-5 in iPSC-CMs) SNVs with significant imbalance per phenotype at FDR<0.05 (**Fig. 3b**), which corresponded to 0.3%-10% of all SNVs tested in each dataset. Shared ASE-SNVs between iPSC-CMs and iPSCs (519 in RNA-Seq and 43 in H3K27ac) showed high concordance of ASE effects (**Fig 3c**) - defined as the mean proportion of the alternate allele across heterozygous sites (Spearman correlation *r* >0.85) - indicating consistency of allelic effects between the two cell types. To confirm that these genetic associations were not spurious, we tested whether the ASE observed in heterozygous individuals was consistent with the overall effect size on the phenotype observed when including homozygous samples. We first calculated the slope (effect size; ß) of the linear relationship between read coverage and genotype for each peak/gene containing an ASE-SNV, and then compared the ASE effect to the corresponding ß in each region (**Fig. 3d-f**, using the most significant ASE-SNV per region). We observed a significant (*P* < 0.05), positive relationship for all molecular phenotypes, with the highest correlation in NKX2-5 peaks (*r*=0.69, Spearman correlation), followed by H3K27ac peaks (*r*=0.57, in iPSC-CMs and *r*=0.61 in iPSCs), and RNA-Seq (*r*=0.40 in iPSC-CMs and *r*=0.47 in iPSCs). These data demonstrate that the majority of allele-specific effects identified in both iPSC-CMs and iPSCs are due to genetic variation, and that, among the three molecular phenotypes examined, NKX2-5 peaks had substantially more ASE-SNVs and showed the highest consistency across individuals.

**Figure 3.**
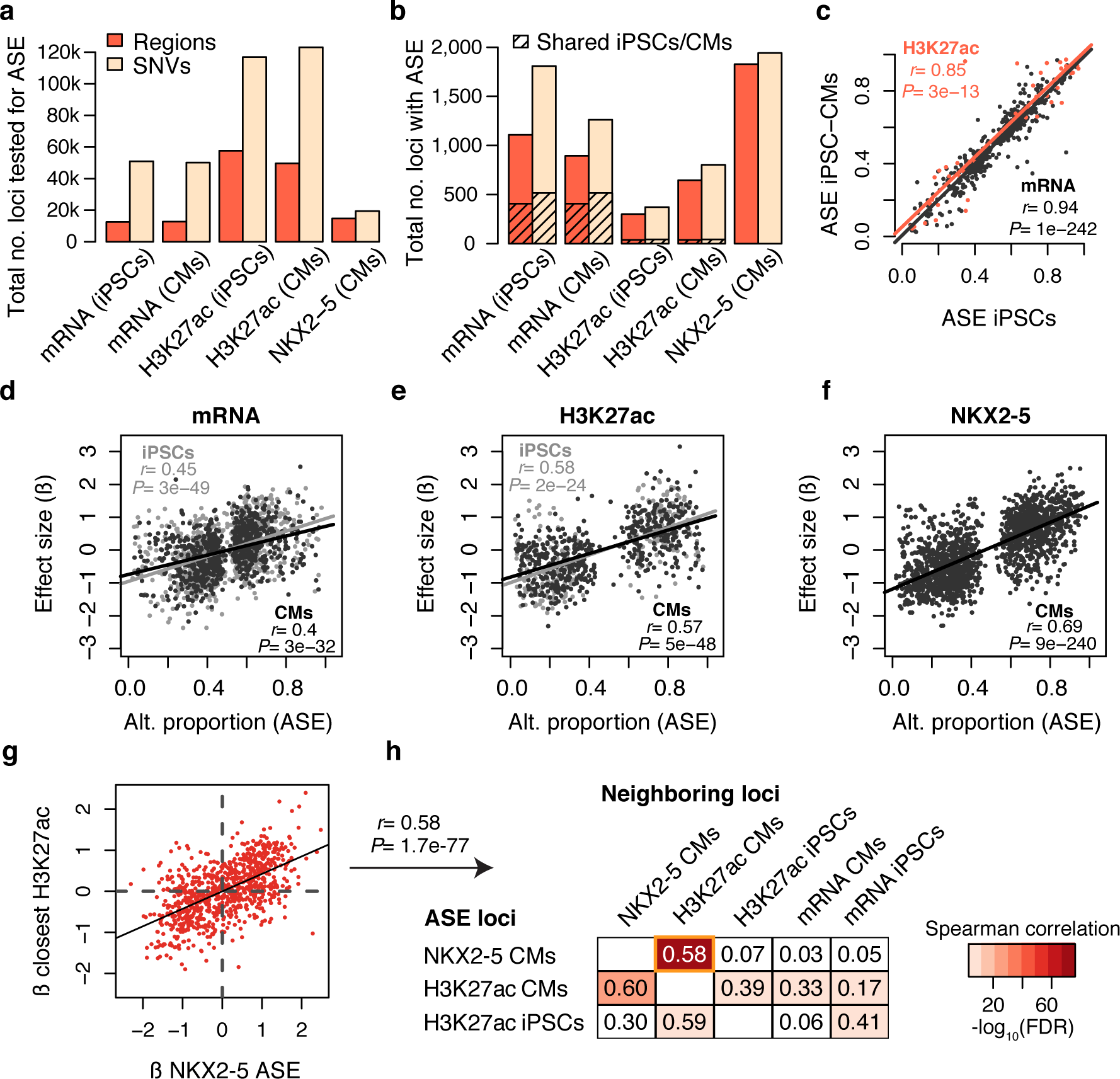
Identification of coordinated allele-specific effects (ASE) in gene expression, histone acetylation and NKX2-5 binding in iPSCs and iPSC-CMs. (**a**) Total number of regions and heterozygous SNVs tested for ASE across all samples in each dataset. (**b**) Total number of heterozygous SNVs and corresponding regions across all individuals with ASE at FDR< 0.05. The number of ASE shared between iPSCs and iPSC-CMs is indicated by hatches. (**c**) Scatterplot of the alternate allele proportion at shared ASE-SNVs between iPSCs and iPSC-CMs for RNA-Seq and H3K27ac. Spearman correlation statistics are indicated. (**d-f**) Scatterplots of the mean proportion of the alternate allele of SNVs with ASE in heterozygous individuals and the effect size of each ASE-SNV, expressed as the slope of linear regression (ß) between gene expression or peak density and genotypes of all seven individuals. Spearman correlation statistics are indicated. (**g**) Scatterplot showing relationship between effect sizes (ß’s) of ASE-SNVs in NKX2-5 peaks on both NKX2-5 and H3K27ac phenotypes. (**h**) Table showing Spearman correlation coefficients of effect sizes between pairs of different molecular phenotypes. Correlations were calculated between ß’s of SNVs that showed ASE in ChIP-Seq datasets (rows) and ß’s of the same variant for the closest gene or peak in a different molecular phenotype dataset (columns). The table is color-coded based on —log_10_ Spearman correlation FDR corrected *P*-values.

### NKX2-5 and H3K27ac allele-specific effects are positively correlated

Genetic loci associated with differential TF binding between individuals often show coordinated effects across different molecular traits^41^. To examine if NKX2-5 loci with ASE were correlated with H3K27ac and gene expression ASEs, we compared the β of ASE-SNVs identified within ChIP-Seq peaks with the effect size of the same SNV on neighboring regions from different molecular phenotypes (nearest peak or nearest gene) (**Fig. 3g-h**). The strongest positive correlation was found between NKX2-5 and H3K27ac genetic effects in iPSC-CMs (Spearman correlation coefficient r=0.58 for NKX2-5 ASE-SNVs, **Fig. 3g**, and *r*=0.60 for H3K27ac ASE-SNVs), supporting the role of NKX2-5 binding in enhancer and promoter activation in these cells. However, genetic effects on NKX2-5 binding were not positively correlated with the expression of neighboring genes (**Fig. 3h**), possibly due to NKX2-5’s dual role as an activator or repressor^31,42^. We also observed that, when iPSC-CMs or iPSCs had H3K27ac ASE, the effect sizes were positively correlated (*r*=0.39, *r*=0.59) with H3K27ac peaks in nearby or overlapping regions in the other cell type, suggesting conserved genetic effects at shared enhancers and promoters. On the other hand, while H3K27ac ASE effect sizes were moderately correlated with gene expression in the corresponding cell type, they were not correlated with gene expression in the other cell type (*r*=0.33 and *r*=0.41 within the same cell type, and *r*=0.17 and *r*=0.06 for mismatched comparisons **Fig. 3h**). These results show that, in both the iPSC-CMs and iPSCs, genetic variation underlies coordinated, and cell-type specific differences across multiple molecular phenotypes; of note, while NKX2-5 and H3K27ac ASE-SNVs were highly correlated, altered NKX2-5 binding was not positively correlated with gene expression changes, consistent with a more complex function as both an activator and repressor.

### Alterations of cardiac TF binding motifs underlie NKX2-5 ASE-SNVs

We next investigated whether genetic variants associated with altered NKX2-5 binding affected sequence motifs of TF binding sites. We selected the most enriched motifs in NKX2-5 peaks calculated using HOMER, which included the NKX2-5 homeobox motif (cognate motif), as well as motifs of other heart development TFs (GATA4, TBX5, TBX20, MEF2A/C and MEIS1, **Supplementary Table 4**) (secondary motifs). For both alleles of all heterozygous SNVs tested for ASE within NKX2-5 peaks, we calculated the motif position weight matrix (PWM) score of each motif. We then compared SNVs with ASE, to SNVs without ASE, and observed that the former was enriched for altered motifs (Fisher’s exact test FDR<0.05) (**Fig. 4a**). Out of the 1,941 NKX2-5 ASE-SNVs, 735 (37.8%) modified at least one of the twelve tested TF motifs: 94 (4.8%) modified both the cognate and a secondary motif, 247 (12.7%) modified only the cognate motif, and 394 (20.3%) modified one or more secondary motifs. Next, we asked whether the preferred allele (highest read count) of each ASE-SNV was associated with a higher predicted motif score. For most motifs (EOMES and CTCF were exceptions), the preferred allele increased the motif score in 70-88% of SNVs (**Fig. 4b**), and the allelic proportion of ASE-SNVs positively correlated with the change in motif score, supporting an underlying causal effect for the majority of these SNVs (**Fig. 4c-d, Supplementary Fig. 5**). We additionally observed that ASE-SNVs tended to affect core, conserved positions within the motif more frequently than they affected less conserved positions (**Fig. 4e-h**), indicating a stronger effect on TF binding affinity. These data indicate that ~40% of sites containing NKX2-5 ASE-SNVs have altered motifs for NKX2-5 and/or for other known cardiac TFs, suggesting that differential allelic binding of NKX2-5 at these sites likely occurred either directly, due to alterations of its own binding sequence, or indirectly, via alterations of TF binding sites of co-binding partners.

**Figure 4.**
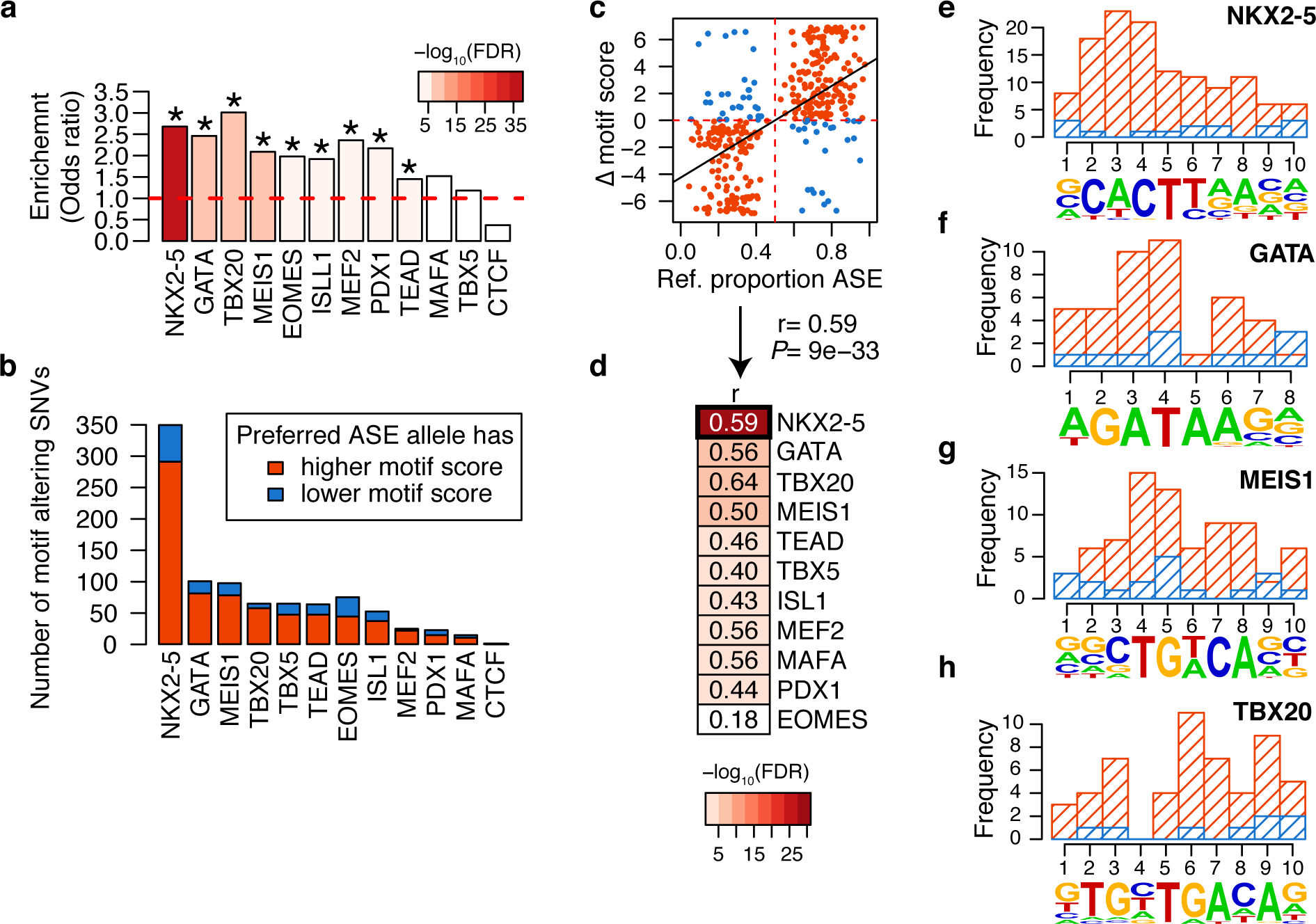
Transcription factor binding motifs are altered by SNVs with ASE in NKX2-5 ChIP-Seq. (**a**) Odds ratios from Fisher’s exact test comparing the proportion of motif-altering SNVs between variants with ASE and variants without ASE in NKX2-5 ChIP-Seq peaks. Bars are color coded according to -log_10_ of FDR corrected P-values. Asterisks indicate enrichment at FDR corrected P-value <0.05. (**b**) Number of TFBS motifs that were strengthened (red) or weakened (blue) by the preferred allele of ASE-SNVs identified in NKX2-5 ChIP-Seq. (**c**) Scatterplot of the reference allele proportion at ASE SNVs and the difference of NKX2-5 motif score between reference and alternate alleles. Spearman correlation coefficient and P-value are indicated at the bottom. Dots are color-coded as in **b**. (**d**) Summary table of Spearman correlation statistics calculated as in **c**, for all motifs tested (see **Supplementary Fig. 5** for the other scatterplots). (**e**-**h**) Frequency of ASE-SNVs altering different positions within the motifs of (**e**) NKX2-5, (**f**) GATA, (**g**) MEIS1 and (**h**) TBX20. NKX2-5, GATA and TBX20 PWMs were obtained using de-novo motif finding. Bars are color-coded as in **b.** Blue bars overlap the red ones (i.e. they are not stacked).

### NKX2-5 ASE-SNVs modulate cardiac-specific gene expression

We next examined if NKX2-5 ASE-SNVs were associated with cardiac-specific effects on gene regulation. We compared the enrichment of NKX2-5 and H3K27ac ASE-SNVs with quantitative trait loci (QTL) from diverse cell types including: DNase hypersensitivity QTLs (dsQTLs) in lymphoblastoid cell lines (LCLs)^43^, expression QTLs (eQTLs) from iPSCs^22^, and eQTLs from 13 combined studies obtained from Haploreg^44^ (“combined tissues”) (**Fig. 5a-c and Supplementary Table 5**). In iPSC-CMs, H3K27ac ASE-SNVs were enriched over SNVs without ASE for all three types of QTLs (OR for dsQTLs = 4.55, for iPSC eQTLs = 1.61, for combined tissues eQTLs = 1.62, *P*<1×10^−4^, Fisher exact test); on the other hand, H3K27ac ASE-SNVs in iPSCs were only enriched for iPSC eQTLs (OR = 2.76, *P*=1.2×10’^9^). Of note, NKX2-5 ASE-SNVs were significantly depleted for iPSC or combined tissue eQTLs (OR = 0.54 and 0.74, *P*<1×10^−6^, respectively), suggesting that they exert regulatory functions only in cardiac tissues.

**Figure 5.**
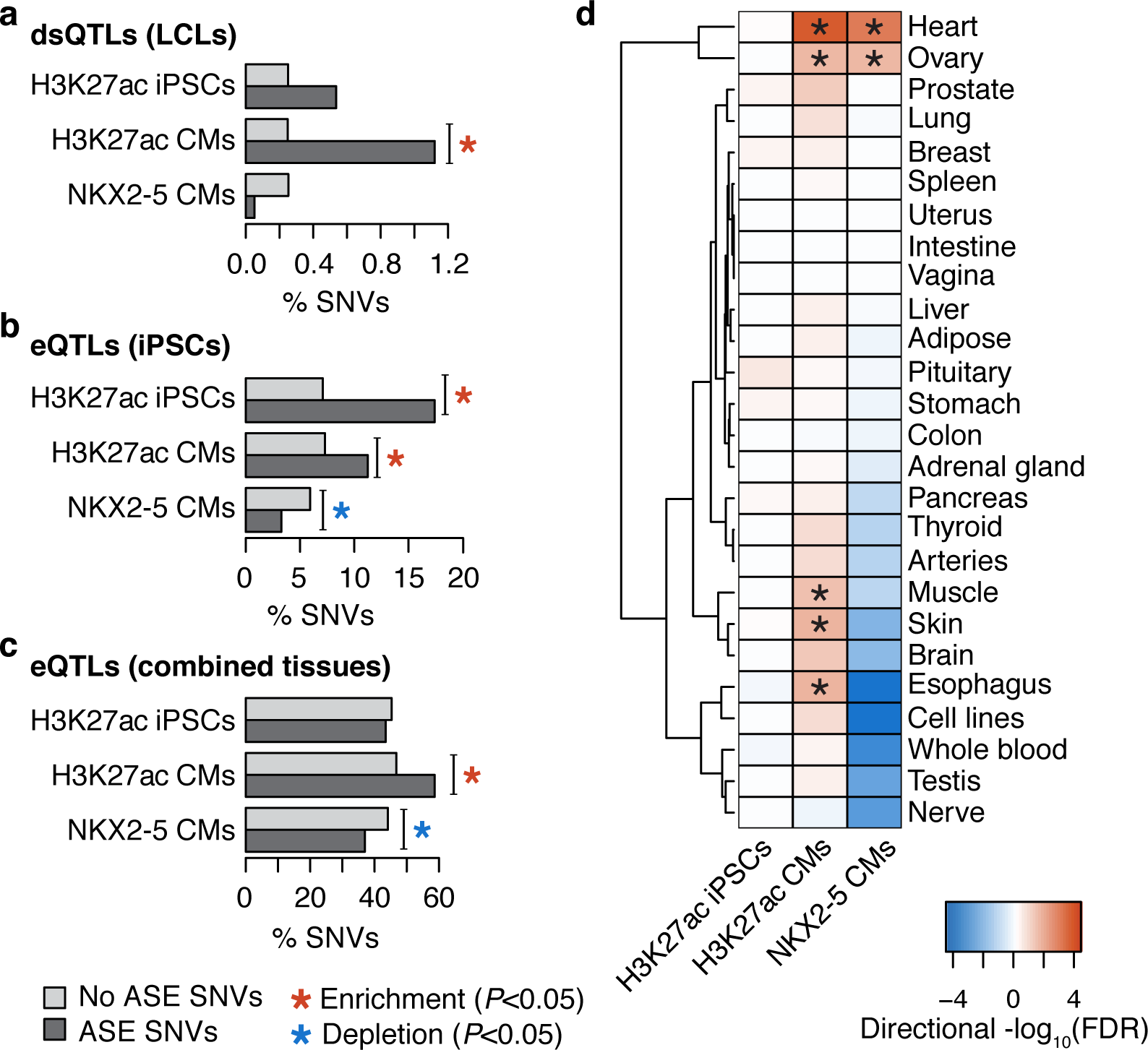
Enrichment of ChIP-Seq ASE variants for known QTLs. (**a-c**) Histograms showing the percentage of SNVs with and without ASE in each ChIP-Seq (H3K27ac in iPSCs and iPSC-CMs and NKX2-5 in iPSC-CMs) and overlapping (**a**) dsQTLs from LCLs^43^, (**b**) eQTLs from iPSCs^22^ and (**c**) combined eQTLs identified in different tissues^44^. Asterisks indicate Fisher’s exact test P-value <0.05 and are colored red or blue for enrichment or depletion, respectively. (**d**) Heatmap showing enrichment of ASE variants for tissue-specific eQTLs^18^ (similar tissues in GTEx merged - see Methods). Asterisks indicate Fisher’s exact test FDR corrected P-value <0.05. Heatmap is colored based on -log_10_ of FDR corrected P-values, with negative sign if odds ratio was <1.

We therefore investigated if NKX2-5 ASE-SNVs were enriched for heart-specific eQTLs. NKX2-5 and H3K27ac ASE-SNVs were compared with SNVs without ASE to assess enrichment for tissue-specific eQTLs (defined in methods) in 26 tissue types from the GTEx project (v6)^18^. ASE-SNVs in both NKX2-5 and H3K27ac peaks in iPSC-CMs were more enriched for heart-specific eQTLs (Fisher exact test: ORs = 1.81 and 2.43, FDRs=1.2×10^−3^ and 2.4×10^−4^, respectively, **Fig. 5d**) than other tissue-specific eQTLs, while H3K27ac ASE-SNVs in iPSCs were not enriched for any GTEx tissue-specific eQTL. Of note, there were 55 NKX2-5 ASE-SNVs that overlapped a heart-specific eQTL, of which 9 affected the NKX2-5 binding motif, and 13 affected one or more of the other cardiac TF motifs in **Fig. 4** (**Supplementary Table 5**). These results indicate that ASE-SNVs in the iPSC-CM lines are enriched for tissue-specific regulatory variants associated with molecular traits in previous studies. Overall, consistent with its importance as a cardiac identity transcriptional regulator, we found that SNVs affecting the binding of NKX2-5 and other cardiac TFs (with which NKX2-5 cooperatively binds) are likely to modulate cardiac-specific gene expression.

### NKX2-5 ASE-SNVs are enriched for GWAS associations with EKG traits

Based on the fact that GWAS variants near the *NKX2-5* gene have been previously associated with EKG traits^6,10,34^, we hypothesized that the altered binding of NKX2-5 in other GWAS loci could be causally implicated in these traits. We first examined if NKX2-5, H3K27ac, or ATAC peaks from iPSC-CMs were enriched for GWAS-SNPs (lead SNPs or SNPs in linkage disequilibrium; LD, r^2^>0.8) for six EKG traits (heart rate, PR interval, QT interval, QRS duration, atrial fibrillation and P-wave duration), compared with GWAS-SNPs from 119 other traits having a comparable number of associated SNPs. We observed a strong relative enrichment for several EKG traits (Binomial test FDR<0.05, **Fig. 6a-c, Supplementary Fig.6**), with QRS duration GWAS-SNPs and heart rate GWAS-SNPs being the top two enriched traits in NKX2-5 peaks. We also examined H3K27ac and DHS from Roadmap cardiac tissues, which similarly showed high enrichment for all EKG GWAS-SNPs, while H3K27ac and DHS from iPSCs did not (Supplementary Fig. 6). These data show that iPSC-CM and Roadmap cardiac tissues have similar enrichment profiles for EKG trait-specific regulatory variants.

**Figure 6.**
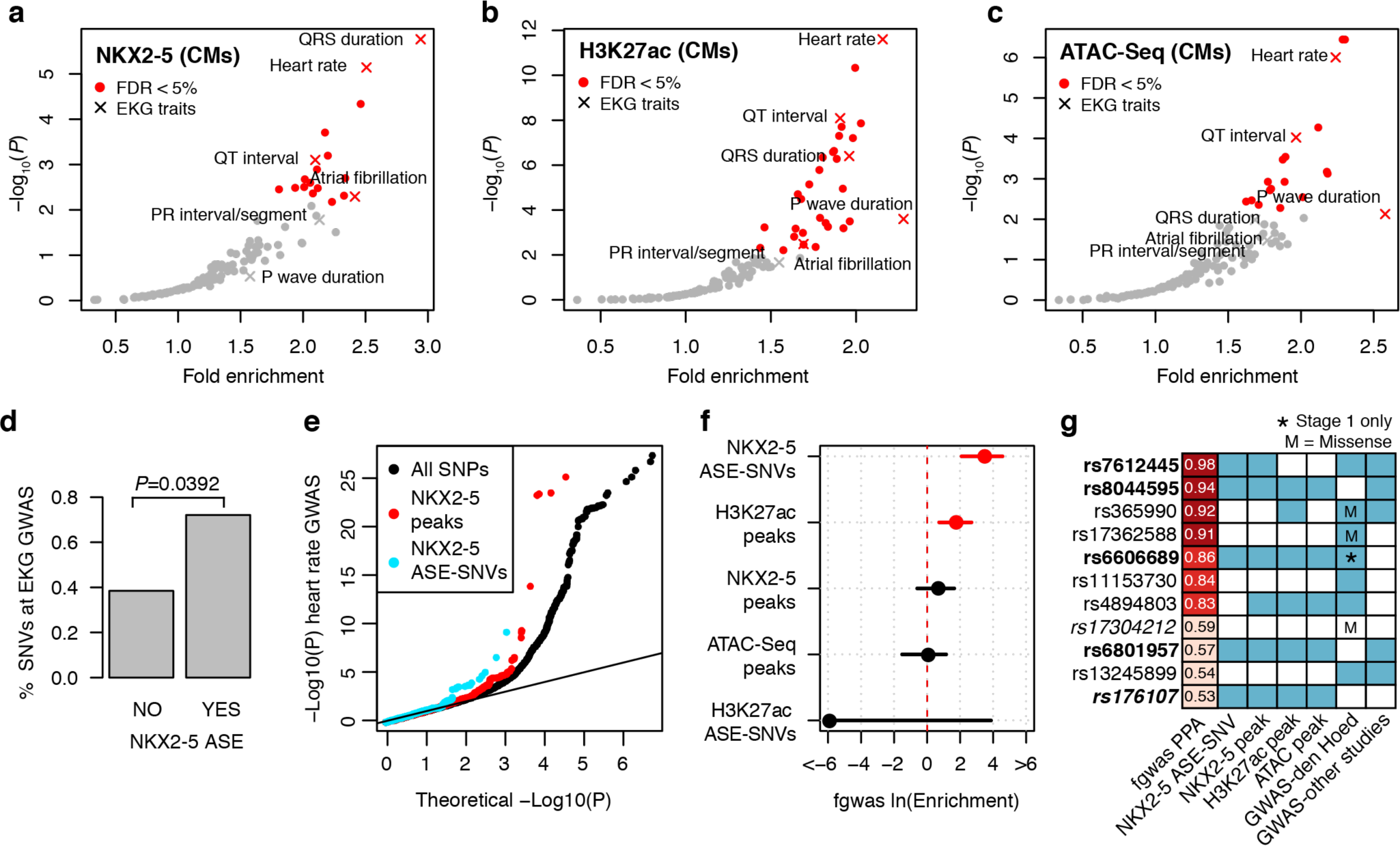
Enrichment of iPSC-CM NKX2-5, H3K27ac and ATAC-Seq peaks in GWAS loci. (**a-b**) Volcano plots showing -log_10_ P-values and fold enrichment for GWAS loci in (**a**) NKX2-5, (**b**) H3K27ac, and (**c**) ATAC-Seq peaks in iPSC-CMs. Red symbols indicate significant enrichment at FDR corrected P-value <0.05. In total 125 gWAs traits were tested, of which 6 were for EKG traits. (**d**) Percentage of SNVs tested for NKX2-5 ASE that overlapped an EKG GWAS SNP and that either did not show or showed ASE. Fisher’s exact test P-value is given. (**e**) Inflation of small P-values from a GWAS meta-analysis for heart rate (den Hoed et al^6^), with enrichment among SNPs in NKX2-5 peaks and NKX2-5 ASE-SNVs. (**f**) Forest plot showing the natural log of the enrichment of heart rate association signal (calculated by fgwas), and 95% confidence intervals, within the genomic annotation listed on the y-axis. Red points denote significantly enriched annotations, and black points denote non-significant enrichments. (**g**) Table showing the SNPs with >0.5 posterior probability of causality (PPA) calculated by fgwas and corresponding annotations conferring positive weights: overlap with NKX2-5 ASE-SNVs, NKX2-5, H3K27ac and ATAC-Seq peaks. SNPs within loci that showed genome-wide significance for heart rate association in the den Hoed et al. GWAS meta-analysis^6^ and/or in other GWAS studies^7^ are indicated. Bolded SNPs denote NKX2-5 ASE SNVs and italicized SNPs denote novel GWAS loci.

To estimate if differential binding of NKX2-5 might have a role in these EKG phenotypes, we examined whether the iPSC-CM NKX2-5 ASE-SNVs were more likely to be EKG GWAS-SNPs than SNVs that did not show ASE. To obtain a sufficient number of GWAS loci containing the variants tested for ASE in these subjects, we combined GWAS-SNPs from all six EKG traits (417 independent SNPs), and intersected them with NKX2-5 peaks. In total, there were 121 GWAS-SNPs within NKX2-5 peaks, of which 81 were heterozygous in the family and had sufficient read coverage to be tested for ASE. Fourteen of these GWAS-SNPs were ASE-SNVs (**Table 1**), which resulted in a significantly higher proportion of EKG GWAS-SNPs among NKX2-5 ASE-SNVs compared to SNVs in NKX2-5 peaks without ASE (Fisher’s exact test, OR=1.88, P=0.0392) (**Fig. 6d**). Among these 14 NKX2-5 ASE-SNVs at EKG GWAS loci, seven (rs590041, rs35176054, rs7612445, rs6606689, rs3807989, rs6801957 and rs10841486) were evolutionary conserved in mammals (SiPhy conservation^44^), and/or altered a cardiac TF motif, (Table 1); and three (rs7612445, rs7986508, rs590041) overlapped heart-specific eQTLs from GTEx. These results suggest a functional link between NKX2-5 binding, cardiac-specific gene expression, and EKG phenotypes at these loci. Based on these findings, we concluded that differential binding of NKX2-5 likely plays a role in EKG phenotypes, and as such, NKX2-5 ASE-SNVs could be used to prioritize candidate causal variants within EKG GWAS loci.

**Table 1.**
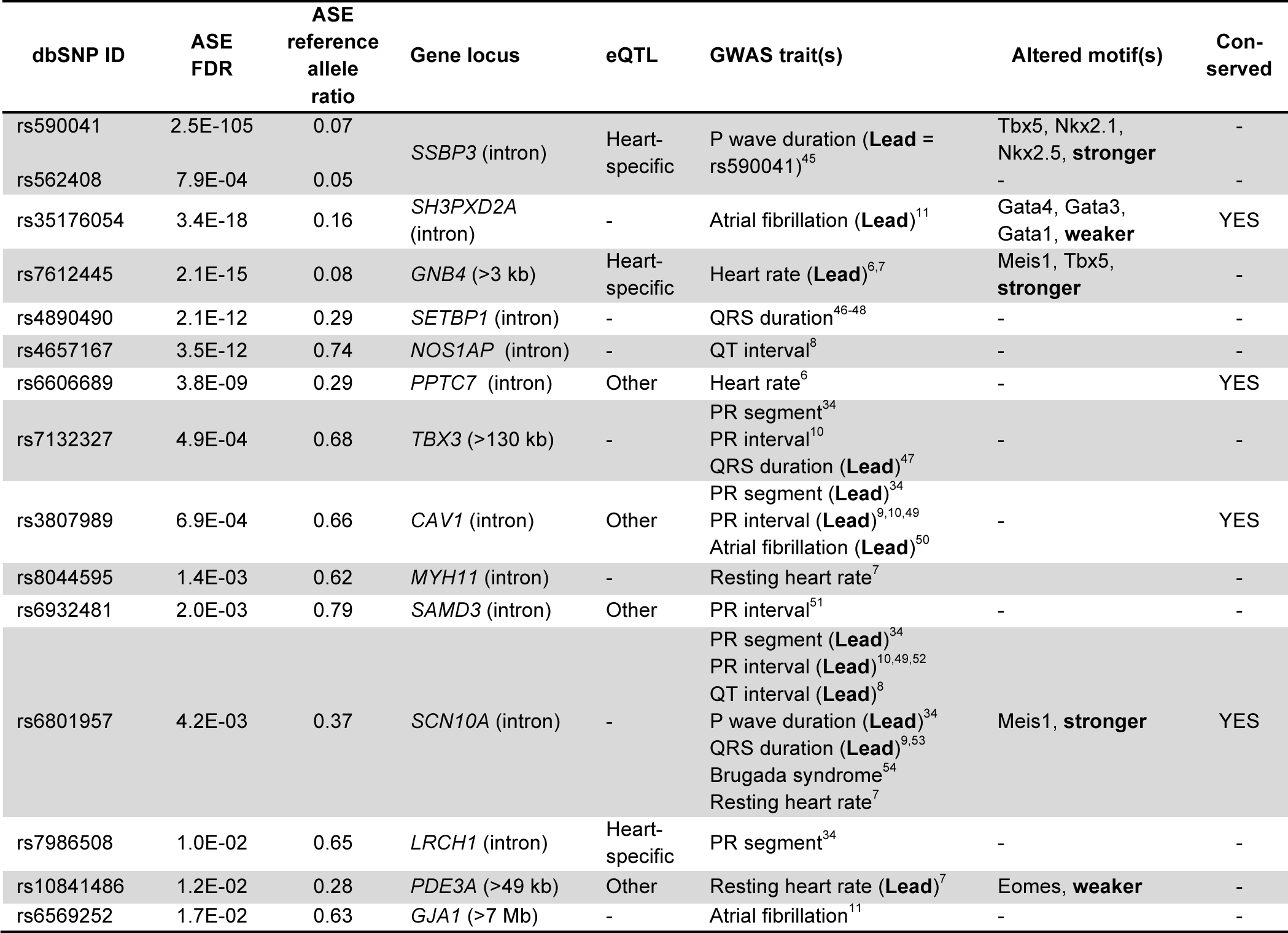
Allelic binding of NKX2-5 at GWAS loci for electrocardiographic (EKG) traits. Fourteen GWAS loci for EKG traits overlapping NKX2-5 ASE-SNVs, ordered by *P*-value for imbalance. For each SNV, we indicate the dbSNP ID (build 137), ASE corrected *P*-value (FDR) combined across heterozygous samples, ASE reference allele ratio, the closest genes and relative location of the SNV, known association with gene expression (eQTL) and in which tissue (heart-specific = restricted to left ventricle and/or atrial appendage in GTEx, other = any other tissue or cell line), associated EKG GWAS traits and if the SNV is the lead variant, altered motifs, if the preferred ASE allele strengthened or weakened the binding motif and conservation in mammals. Additional annotations are reported in **Supplementary Table 5.**

### NKX2-5 ASE-SNVs prioritize causal variants within heart rate GWAS loci

To examine the extent to which NKX2-5 ASE could be used to prioritize causal variants, we utilized fgwas, a statistical framework that integrates functional genomics annotations and GWAS data to identify putative causal variants as well as potentially novel loci55. We focused on a GWAS meta-analysis for heart rate^6^, and performed fine-mapping using summary statistics from this study and the molecular phenotype data (NKX2-5, H3K27ac, and ATAC-Seq) from iPSC-CMs. We observed an enrichment of low P-values from the heart rate GWAS meta-analysis at NKX2-5 ASE-SNVs above the enrichment at all SNVs in NKX2-5 peaks (**Fig. 6e**). Therefore, we treated ASE-SNVs as a separate annotation for fine-mapping, creating 5 possible annotations for fgwas: overlapping an 1) ATAC-seq peak, 2) H3K27ac peak, 3) NKX2 peak, or being an 4) H3K27ac ASE-SNV, 5) NKX2-5 ASE-SNV. We simultaneously quantified the enrichment of heart rate association signal for each annotation genome-wide using fgwas, and found NKX2-5 ASE-SNVs to be the most significantly enriched, followed by H3K27AC peaks (**Fig. 6f**). Next, using fgwas, we refined these enrichment estimates using 10-fold cross-validation; these refined values were used as priors to update the probability for a variant to be causal (posterior probability of association, PPA) within consecutive 1Mb windows across the genome. Out of ~2.5 million tested variants, we found 21 variants with greater than 30% probability of being causal, of which seven were NKX2-5 ASE-SNVs (**Supplementary Table 6**); these findings suggest that altered binding of NKX2-5 accounts for a considerable fraction of the genome-wide genetic contribution underlying variable heart rate. Out of the 11 most significant variants (PPA > 0.50) (**Fig. 6g**), five were NKX2-5 ASE-SNVs; four of these were in loci previously associated with heart rate either in the same meta-analysis^6^ (rs7612445 and rs6606689) and/or in a different GWAS (rs8044595 and rs6801957^7^), and details for these variants are shown in Table 1. While **rs6801957**, identified with a 57% probability of causality (**Fig. 7a**), did not reach genome-wide significance in the heart-rate meta-analysis, it was associated in a larger heart-rate GWAS^7^, as well as in several previous GWAS with other EKG traits^8–10,34,49,52,53^. Moreover, rs6801957 has been experimentally validated as a causal variant for EKG traits. While we predicted that rs6801957 altered a T-box binding sequence and resulted in differential co-binding of NKX2-5, previous functional experiments showed that this variant affects binding of TBX3 and TBX5 and expression of *SCN5A*, the main cardiac sodium channel^56,57^. Overall, these results show that fine-mapping statistical approaches, combined with molecular phenotype data (NKX2-5, H3K27ac, and ATAC-Seq) from iPSC-CMs, can be used to prioritize putative causal variants in heart rate GWAS loci, and suggest that differential NKX2-5 binding underlies the mechanism of numerous heart rate loci across the genome.

**Figure 7.**
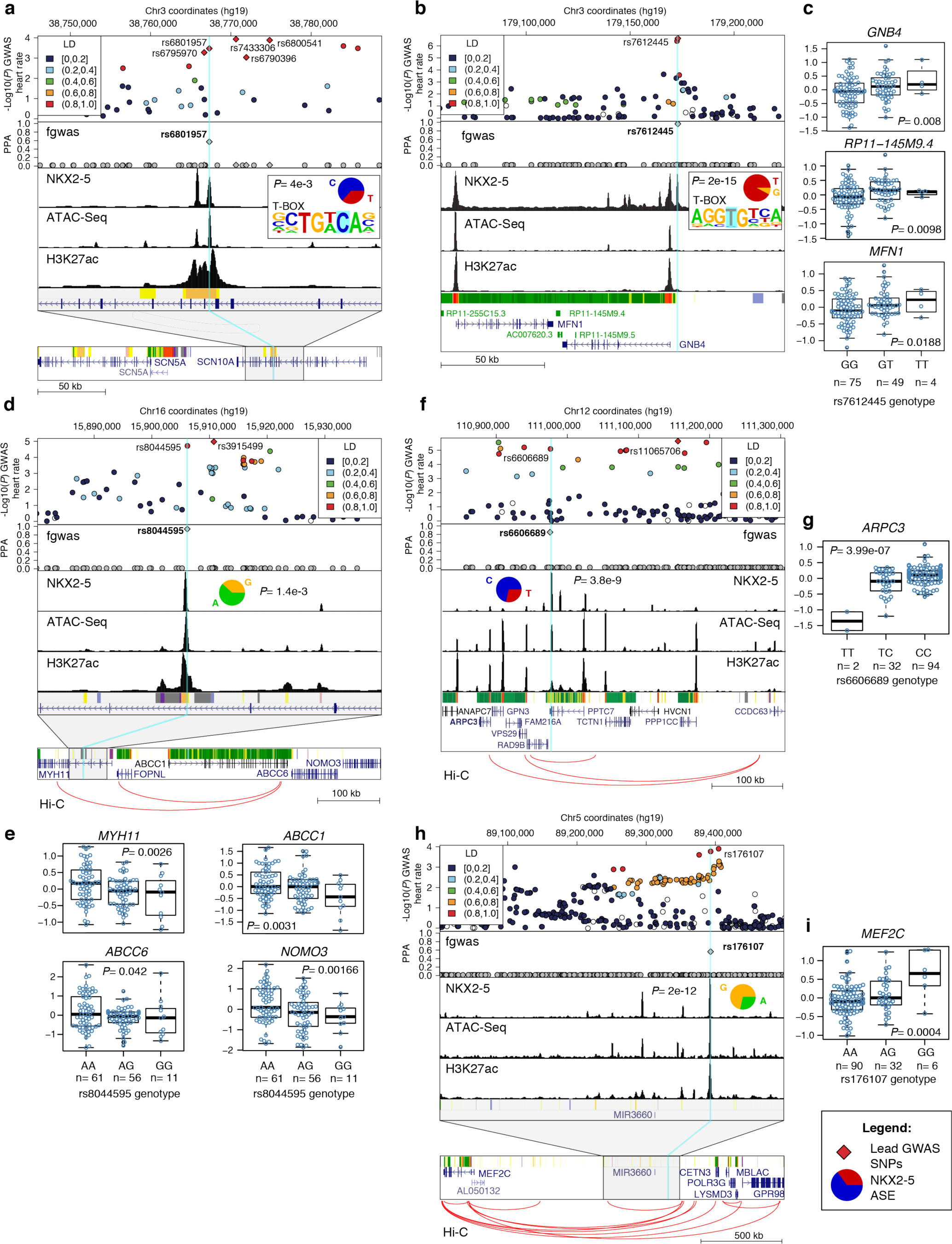
Functional characterization of candidate causal variants at loci associated with heart rate. For each of the five loci (**a, b, d, f** and **h**) the top panel shows the regional plot of association P-values with heart rate^6^; SNPs are color coded based on r^2^ values from the 1,000 Genome Project CEU population^59^; lead GWAS variants in the locus are indicated by a diamond. The second panel shows the posterior probability of causality (PPA) of the variants in the locus calculated using fgwas, and panels three through five show epigenetic tracks from the iPSC-CM metasamples (NKX2-5, ATAC-Seq and H3K27a). The bottom panel shows the Roadmap fetal heart ChromHMM and genes from UCSC genome browser (conventional ChromHMM color code). For **a, d** and **h**, the bottom panel shows the locus at lower scale. For **d, f** and **h**, the locations of Hi-C loops from iPSC-CM are shown in red. For the candidate causal variants (turquoise lines), the allelic imbalance (pie chart) of NKX2-5 ASE and FRD-corrected P-values are shown; for **a** and **b**, the altered TF motif is shown. Significant associations (P <0.05, linear regression) between putative variant genotypes and gene expression of candidate genes in 128 iPSC-CMs from iPSCORE are shown (**c, e, g** and **i**).

### NKX2-5-mediated mechanisms of association with heart rate

To further investigate the mechanisms of association between heart rate and the NKX2-5 ASE-SNVs identified as candidate causal variants, we followed up the three loci previously associated with heart rate (without validated causal variants), and the novel locus with additional experimental data. These data included Hi-C chromatin conformation maps from the same iPSC-CM lines used to generate the other molecular phenotypes in this study^58^ (**Supplementary Table 2A**), in order to identify long-range interactions between the candidate causal variants and target genes, and RNA-Seq data from iPSC-CMs from an additional 128 whole-genome-sequenced subjects^36^, to evaluate the association of NKX2-5 ASE-SNVs genotypes with the expression of candidate genes.

**rs7612445** was previously identified as a lead heart rate GWAS variant^6,7^, as well as a heart-specific eQTL in GTEx for *GNB4* and two non-coding genes *(RP11-145M9.4* and AC007620.3). Our fine-mapping analysis showed rs7612445 had a probability of 98% of being causal, which is further supported by the fact that the variant creates a T-box binding sequence, resulting in increased NKX2-5 binding in the iPSC-CMs (**Fig. 7b**). Analysis of the RNA-Seq data from the 128 iPSC-CM lines showed associations between rs7612445 and expression of *GNB4* and *RP11-145M9.4* as observed in GTEx (**Fig. 7c**), as well as other nearby genes: *MFN1, NDUFB5, ZNF639, PIK3CA*, and *RP11-255C15.4.* For four genes of these genes *(GNB4, RP11-145M9.4, MFN1* and *NDUFB5,)*, the allele of rs7612445 with preferential TF binding was associated with increased gene expression (positively associated, i.e activation), while for the other three genes *(ZNF639, PIK3CA*, and *RP11-255C15.4)*, it was associated with decreased gene expression (negatively associated, i.e. repression). Although it is unclear which gene(s) in the interval are responsible for the GWAS association, the mitochondrial fusion protein *MFN1* was previously indicated as a promising candidate, based on a reduction in heart rate when the gene was knocked-down in both Zebrafish and Drosophila models^6^.

**rs8044595** had a high probability of being causal (94%) in our fine-mapping analysis, but was not detected in the heart rate GWAS meta-analysis for which we used summary statistics^6^. However, it was associated with heart rate in a larger GWAS^7^, with rs8044595 being in perfect LD with the lead SNP (rs3915499) (**Fig. 7d**). Using the Hi-C data, we determined that rs8044595 was in a chromosome loop that would result in contact of the regulatory element with five genes *(MYH11, FOPNL, ABCC1, ABCC6* and *NOMO3)* spanning ~500kb. Gene expression analysis in the 128 iPSC-CMs revealed positive associations with *MYH11, ABCC1, ABCC6* and *NOMO3* (**Fig. 7e**). Among the four genes, *NOMO3* is the most plausible candidate, as the protein is highly expressed in cardiomyocytes^60^, is part of Nodal signaling, which plays an essential role in development, and has been implicated in with congenital heart defects through copy number alterations in the interval encoding the gene ^61,62^.

**rs6606689** had an 86% probability of causality, and was significantly associated through LD with a lead variant (rs11065706) in the first stage of the heart rate GWAS meta-analysis (**Fig. 7f**); however, it did not replicate in stage 2. Among the 10 genes in the same Hi-C loop (~400kb), we found that *ARPC3* was the only gene whose expression was associated with rs6606689 in iPSC-CMs (positive association observed) (**Fig. 7g**). In GTEx, rs6606689 is an eQTL for *ARPC3* in skeletal muscle. These data combined with the fact that actin-related protein 2/3 complex subunit 3 (ARPC3) is implicated in the control of actin cytoskeleton polymerization in cells, supports our fine-mapping analysis suggesting that rs6606689 is likely a causal variant associated with heart rate, potentially by its regulation of *ARPC3.*

**rs176107**, while having a 53% probability of causality, was never reported as either a GWAS variant for any EKG trait, or an eQTL for any tissue; however, it had a P-value of 1.7×10^−4^ for association in the heart rate GWAS meta-analysis, and therefore was considered a “sub-threshold” signal (**Fig. 7h**). Using Hi-C, we determined that the *MEF2C* gene, located ~1.2 Mb away, was brought into close proximity with the NKX2-5 binding site through a long-range interaction. Further, expression analysis of *MEF2C* showed a positive association with rs176107 (**Fig. 7h**). MEF2C is a transcription factor critical for muscle, heart and neuronal development, it interacts with NKX2-5 itself^63^; and a KO in mice has been shown to result in early development lethality due to severe cardiovascular defects^64,65^. Thus, although MEF2C has not previously been linked to cardiac defects in humans, it is as is an excellent candidate gene for heart phenotypes.

Overall these data provide novel functional links between NKX2-5 binding, cardiac gene expression regulation and heart rate variability both at novel and previously identified GWAS loci.

## DISCUSSION

In this study, we showed that characterizing genetic variation that alters the binding of NKX2-5 in human cardiomyocytes could reveal the molecular effects of known EKG GWAS variants, as well as enable the identification of novel variants associated with these cardiac traits. Additionally, we showed the utility of iPSC-derived cardiomyocytes (iPSC-CMs) for interrogating the function of cardiac regulatory variants. Despite the fact that current differentiation protocols result in heterogeneous cell populations that vary across cell lines^25,66^, we have shown that genetic background is a major contributor to molecular phenotype differences in iPSC-CMs, supporting their use for identifying and functionally characterizing cardiac regulatory variants. We leveraged the unique advantages of using iPSC-CMs as a model system, deriving lines from seven whole-genome sequenced individuals in quantities sufficient for conducting RNA-Seq, ChIP-Seq of NKX2-5 and H3K27ac, ATAC-Seq and Hi-C data on the same samples. We utilized these data to conduct functional genomic and fine-mapping analyses in order to study in depth the effects of genetic variants that alter NKX2-5 binding, in a manner that would not have been possible using postmortem tissues.

Within ~38,000 NKX2-5 binding sites, we identified 1,941 genetic variants that altered the binding of the transcription factor. Because we investigated seven individuals in a three-generational family, the statistical power for identifying ASE-SNVs was increased, as there were multiple replicates of allelic imbalance at the same heterozygous SNV. However, we anticipate that analyzing a larger sample size would identify a greater fraction of the NKX2-5 sites affected by genetic variants. For the NKX2-5 sites with differential binding, ~40% had genetic variants that altered the cognate TF motif, or motifs of functionally related cardiac TFs. This indicates that a large fraction of the observed allelic binding of NKX2-5 was either a direct consequence of the SNV, or an indirect consequence resulting from the differential binding of a known co-factor. Combinatorial interactions between key cardiac TFs is known to be an important mechanism for orchestrating the cardiac gene expression program during development^28–30,32^. While natural genetic variation has been shown to affect collaborative binding of lineage determining TFs in mice^67^, our study is the first that we are aware of to show that genetic variants affects these processes in humans, highlighting the power of iPSC derived cell types to study genome-wide how genetic variants affect cooperative binding of key developmental TF complexes in humans.

Coding mutations in, and non-coding variants near *NKX2-5* have respectively been associated with congenital heart defects^33^ and EKG traits^6,10,34^, implicating this TF in a range of cardiac disease in both development and adult stages. Here, our analysis of genome-wide NKX2-5 binding enabled us to investigate its role in cardiac phenotypes through a different genetic mechanism, i.e. variation in TF binding sites resulting in differential expression of target genes. We showed that differential NKX2-5 binding was positively correlated with H3K27ac peaks at iPSC-CM enhancers, but not iPSC enhancers, suggesting that NKX2-5 ASE-SNVs altered cardiac specific enhancer activity. These findings are consistent with the fact that we found enrichment for GTEx heart-specific eQTLs in both NKX2-5 and H3K27ac ASE-SNVs in iPSC-CMs. Importantly, out of all the molecular phenotypes examined, NKX2-5 ASE-SNVs were the more strongly associated with EKG loci, thereby implicating NKX2-5 in the development of these traits, and indicating that NKX2-5 ASE-SNVs could be used to prioritize putative causal variants. Analyzing a heart rate meta-analysis using a fine-mapping method that integrates GWAS summary statistics with functional annotations revealed four NKX2-5 ASE-SNVs with a high probability of causality at known loci, and one NKX2-5 ASE-SNV at a novel locus with a lower but still substantial probability of being causal. As a proof that this approach was effective to prioritize causal variants, one of the NKX2-5 ASE-SNVs (rs6801957) had been previously investigated in detail and had been shown to be functionally implicated in the association with heart rate and other EKG traits based on several experimental evidences including differential TBX3/TBX5 binding, reporter assays and association with expression of SCN5A^56,57^.

Further investigation of NKX2-5 heart rate loci using Hi-C generated from the same iPSC-CMs, and gene expression in iPSC-CMs derived from 128 individuals revealed an association between the putative causal NKX2-5 ASE-SNVs and expression of nearby or distal candidate target genes. Notably, *MEF2C* has not been previously associated in genetic studies with cardiac traits, but based on its involvement in cardiac morphogenesis, it is likely that its altered expression would result in cardiac phenotypes. *MEF2C* may not have previously been identified as an eGene because it is located ~1.2 Mb from rs176107, and eQTL studies typically only examine SNPs and genes within 1 Mb of each other for associations^18^; thus, these data highlight the importance of using 3D regulatory maps for eQTL studies. Our findings provide plausible causal mechanisms underlying associations between differential NKX2-5 binding and heart rate at four previously established and one novel locus. For these reasons, the NKX2-5 ASE-SNVs identified in this study as overlapping known and novel EKG loci, and the genes with associated expression, are promising targets for future experimental studies.

Finally, our study demonstrates that analyzing the allelic binding of master developmental TFs in iPSC-derived cell types is highly effective to identify genetic variation important for disease; and suggests that expanding this approach to study other cardiac TF (such as TBX5, GATA4 and MEF2C) in larger sample sizes could potentially identify and characterize many of the regulatory variants that play a role in cardiac traits and diseases.

## METHODS

**Subjects and iPSC derivation**. We selected seven individuals of Asian and European descent in the iPSCORE resource^36^ (**Supplementary Table 1**) that are part of a three-generational family. Our study design included three genetically unrelated subjects and two parent-offspring quartets, which enabled us to examine the inheritance of genetic effects. Fibroblasts from skin biopsies of each subject were reprogramme d using) non-integrative Sendai virus^68^ and genomic integrity and pluripotency of iPSCs was assessed at passages 12-13 as described in Panopulos et al.^36^. Briefly, genomic DNA from iPSCs and peripheral blood were analyzed using Illumina HumanCoreExome arrays and Genome Studio. CNVs present in the iPSCs but not in blood DNA were detected both visually (using B-allele frequencies plots for each chromosome) and automatically (using the Nexus Copy Number software; BioDiscovery). For pluripotency analysis, live iPSCs were stained with fluorescent antibodies (Biolegend) against the pluripotency markers SSEA4 and TRA-1-81^69^ as well as isotope controls and analyzed by flow cytometry. For 5 individuals, we analyzed one iPSC line (“clone”) and for two individuals we analyzed two iPSC lines (**Fig. 1**). The nine iPSC lines were harvested in multiple replicates between passages 12 to 40; a total of 35 different iPSC harvests were used in this study (**Supplementary Table 2**). All subjects provided written informed consent for participation to this study. This study was approved by the Institutional Review Boards of the University of California at San Diego (Project #110776ZF).

**Differentiation of iPSCs into cardiomyocytes**. The nine iPSCs were differentiated into cardiomyocytes (iPSC-CMs) in multiple independent differentiations, according to the monolayer protocol described by Lian et al.^37^, resulting in a total of 26 iPSC-CM samples (biological replicates; **Supplementary Table 2**). Twelve of the iPSC-CM samples were subjected to selection using 4mM Sodium L-Lactate media^70^ following dissociation and re-plating at day 15, and collected at day 25. Fourteen of iPSC-CM samples were collected at day 15, of which one was subjected to lactate purification at day 11. At the day of collection, iPSC-CMs from three to four T150 flasks were dissociated using Accutase (Thermo Scientific), pooled, counted and separated into different aliquots. About 6 × 10^7^ cells were fixed with formaldehyde and frozen for ChIP-Seq. Cells (2 × 10^7^) were lysed and stored in RLT plus buffer (Qiagen) for RNA extraction. Nuclei from 2 × 10^5^ cells were frozen for ATAC-Seq. The same protocols of dissociation and collection of samples for RNA-Seq, ChIP-Seq and ATAC-Seq were applied to non-differentiated iPSC lines. Not all of the data types were obtained for each cell line replicate, due to lack of material availability or poor data quality.

**Characterization of cardiomyocyte cellular markers**. For flow cytometry, 1 × 10^6^ fixed iPSC-CMs were permeabilized and blocked for 30 min at room temperature in 0.5% BSA, 5% goat serum, 0.2% TX-100 in PBS and stained for the cardiac marker cardiac troponin T (TNNT2) (Thermo Scientific MA5-12960) for 30 min, followed by anti-mouse Alexa Fluor 488 secondary antibody (Life Technologies A11001) for 30 min. Washes, incubation with antibodies, and flow cytometry analysis were done in 0.5% BSA in PBS. Cells were acquired on a FACSCanto system (BD Biosciences) and analyzed using FlowJo.

For immunofluorescence, three iPSC-CM lines were fixed using 4% PFA in PBS, blocked and permeabilized for 1 h at 37°C with 5% BSA, 5% serum, 0.1% Triton X-100 in PBS. Cells were then incubated with 1:200 dilutions of primary antibodies against TNNT2 (Thermo Scientific MA5-12960) and MYL7 (Synaptics Systems 311011) overnight at 4°C. Slides were washed three times with PBS and incubated for 2 h at room temperature with secondary antibody (AlexaFluor 488) and 20 min with DAPI. Cells were washed with 1× PBS and mounted using Vectashield. Sections were analyzed using a Leica SP5 confocal microscope.

**Whole-Genome Sequencing**. Genomic DNA from the 7 individuals was whole genome sequenced as a part of the iPSCORE collection^22^. Briefly, DNA was isolated from peripheral blood using DNEasy Blood & Tissue Kit (Qiagen), quantified, normalized, and sheared with a Covaris LE220 instrument. Illumina (TruSeq Nano DNA HT kit) DNA libraries were prepared, quality checked and sequenced on a HiSeqX (150 base paired-end). Reads were aligned against human genome b37 with decoy sequences^71^ using BWA-MEM and default parameters^72^. The resulting BAM files were sorted using Sambamba^73^ and duplicate reads were marked using biobambam2^74^. Variant calling was performed using the GATK best-practices pipeline^75,76^ on BAM files separated into individual chromosomes. We performed quality control for the genotypes of SNVs and indels using GATK’s Variant Quality Score Recalibration (VQSR) and used a sensitivity cutoff of 99.5% for SNVs and 99% for indels^77^.

**RNA-Seq**. Total RNA was isolated using the Qiagen RNAeasy Mini Kit from frozen RTL plus pellets, including on-column DNAse treatment step. RNA was eluted in 60 μl RNAse-free water and run on a Bioanalyzer (Agilent) to determine integrity. Concentration was measured by Nanodrop. Illumina Truseq Stranded mRNA libraries were prepared and sequenced on HiSeq2500, to an average of 40 M 100 bp paired-end reads per sample. RNA-Seq reads were aligned using STAR^78^ with a splice junction database built from the Gencode v19 gene annotation^79^. Gene-based expression values were quantified using the RSEM package (1.2.20)^80^ and normalized to transcript per million bp (TPM).

**ChIP-Seq**. For histone modification H3K27ac, 2 × 10^6^ fixed cells were lysed in 60 μl of MAGnify™ Chromatin Immunoprecipitation System Lysis Buffer (Thermo Scientific) and sonicated using Bioruptor 200 (Diagenode) for 35-45 min of 30 sec on/30 sec off cycles. H3K27ac antibodies (Abcam ab4729, lots GR183922-2 (1.75 μg) or GR184333-2 (1 μg)) were coupled for 2 hours to ProteinG Dynabeads (Thermo Scientific), and used for overnight chromatin immunoprecipitation in IP buffer (1% Triton-X, 0.1% DOC, 1x TE, 1x Roche Complete Proteinase Inhibitor tablets (RCPI)). Beads were washed five times with washing buffer (50 mM Hepes pH 8, 1% NP-40, 0.7% DOC, 0.5M LiCl, 1mM EDTA and 1x RCPI) and once with TE buffer. For NKX2-5, 1-2 × 10^7^ cells were lysed in 300 μl RIPA buffer (1×PBS, 1% NP-40, 0.5% DOC, 0.1% SDS, RCPI) and sonicated for 70-80 min with instrument and setting as above. Five μg of NKX2-5 antibody (Santa Cruz Biotechnology, sc-8697x, lot C0113) were incubated with Dynabeads for 2 hours and washed with BSA 0.5% in PBS. Chromatin was diluted to 1 ml of RIPA buffer and added to the beads for overnight IP. Five washes were performed with washing buffer (50 mM Hepes pH 8, 1% NP-40, 0.7% DOC, 0.5M LiCl, 1mM EDTA and 1x RCPI), followed by one wash with TE. DNA was eluted and reverse crosslinked overnight in elution buffer (10 mM Tris-HCl pH 8, 1 mM EDTA, 1% SDS) at 65°C. DNA was purified using Qiagen MinElute PCR Purification kit, quantified by Qubit (Thermo Scientific) and submitted to library preparation and barcoding using KAPA Hyper Library preparation kit (KAPA Biosystems). Libraries were sequenced on an Illumina HiSeq2500 or a HiSeq4000 to an average of 35 M 100 bp paired-end reads per sample.

ChIP-Seq reads were mapped to the hg19 reference using BWA^72^. Duplicate reads, reads mapping to blacklisted regions from ENCODE, reads mapping in chromosome other than chr1-chr22, chrX, chrY, and read-pairs with mapping quality Q<30 were filtered. Peak calling was performed using MACS2 (‘macs2 callpeak -f BAMPE -g hs -B –SPMR –verbose 3 –cutoff-analysis –call-summits -q 0.01’)^81^ with reads derived from sonicated chromatin not subjected to IP (i.e. input chromatin) from a pool of samples used as negative control. Peak coordinates were called from a meta-sample of either iPSCs or iPSC-CMs, generated by pooling the BAM files of each data type across all samples of the given cell type. Quantification of the signal at peaks in each sample was performed using featureCounts^82^. Genome browser screenshots showing data quality are presented in **Supplementary Fig. 2**. Motif enrichment analysis was performed using HOMER ‘findMotifsGenome.pl’^83^ and, for NKX2-5, also using MEME ChIP^84^. Motif analysis of the NKX2-5 ChIP-Seq confirmed a significant enrichment (binomial test, q-value <0.0001) for the NKX2-5 homeobox motif, as well as for the motifs of other heart development TFs (GATA4, TBX5, TBX20, MEF2A/C and MEIS1, **Supplementary Table 4**). Peaks from 14 non-purified and one purified iPSC-CM line showed similar motif enrichment results. Additionally, iPSC and iPSC-CM H3K27ac ChIP-Seq and ATAC-Seq data showed enrichment of stem cell TFs (OCT4, NANOG and SOX family) and cardiac TFs, respectively (**Supplementary Table 4**).

**ATAC-Seq**. The ATAC-Seq protocol has been adapted from Buenrostro et al.^85^. Frozen nuclear pellets of 5 × 10^4^ cells each were thawed on ice, suspended in 50 μL transposition reaction mix (2.5 μL Tn5 transposase in 1x TD buffer, Illumina Cat# FC-121-1030), and incubated for 30 min at 37°C. Reactions were purified using Qiagen MinElute kit, eluted in 10 μL water and amplified using the KAPA real-time library amplification kit (KAPA Biosystems) with barcoded adaptors. PCR reactions were terminated after 10 to 13 cycles and purified using AmPure XP beads (Beckman Coulter). Samples were size selected using SPRIselect beads (Beckman Coulter) to a size range of 150 to 850 kbp and sequenced on an Illumina HiSeq2500 to an average depth of 30 M 100 bp paired end reads.

ATAC-Seq reads were aligned using STAR to hg19 and filtered using the same protocol as for ChIP-Seq. In addition, to restrict the analysis to regions spanning only one nucleosome, we required an insert size no larger than 140 bp, as we observed that this improved sensitivity to call peaks and reduced noise. Peak calling was performed using MACS2 on merged BAM files of iPSC and iPSC-CM meta samples with the command ‘macs2 callpeak –nomodel –nolambda –keep-dup all –call-summits -f BAMPE -g hs -q 0.01’. Genome browser screenshots showing data quality are in **Supplementary Fig. 2**.

**Analysis of gene expression differences between iPSCs and iPSC-CMs**. A matrix of raw gene expression values from 64 RNA-Seq samples (29 iPSCs, 27 iPSC-CMs and 8 RNA-Seq samples from Roadmap including H1-hESC (SRR2453359), HUES64 (SRR2453365), iPS-20b (SRR2453353), iPS-18 (SRR2453345), Right Atrium (SRR578651), Right Ventricle (SRR577591), Left Ventricle (SRR577587), and Fetal Heart (SRR3192433)) was created from the RSEM expected counts, filtered for >1 TPM on average samples, and rounded to integer values. After filtering, 15,725 genes remained from the initial 57,820. All RNA samples were processed as described above in the RNA-Seq section. Expression values were normalized using variance stabilizing transformation *(vst)* implemented in DESeq2^86^. Hierarchical clustering and the heatmap in **Supplementary Fig. 1** were generated using vst-normalized read counts for a panel of 61 selected genes (23 pluripotency genes^38^, 22 regulators of cardiac differentiation factors and 16 cardiomyocyte structure factors^39^) using ‘pheatmap’ package in R^87^. Analysis and plotting of principal components of all 15,725 genes were performed in R (**Fig. 1**).

To identify differentially expressed genes (DEGs) between iPSCs and iPSC-CMs we used a matrix of raw expression counts from the 56 RNA-Seq (29 iPSCs and 27 iPSC-CMs), filtered for average TPM >1 (22,447 genes), and applied DESeq2 with default settings to identify DEGs more than 2-fold and at a BH (Benjamini & Hochberg) FDR of 5%.

Analysis of ChIP-Seq and ATAC-Seq coverage at transcription start site (TSS) regions. We used HOMER^83^ to quantify the read density of ATAC-Seq and ChIP-Seq meta-samples at the TSS of differentially expressed genes between iPSCs and iPSC-CMs (2,444 genes upregulated in iPSCs and 2,863 in iPSC-CMs). For each meta-sample, the command ‘annotatePeaks.pl tss hg19 -list -size 2000 -hist 25 -ghist -d [alignment files]’, was used to obtain the coverage of each meta-sample normalized to 10 M reads, in 25-bp sequential bins across a 2-kb window centered on the TSS. The resulting table of normalized counts was ordered by decreasing order of log_2_ ratio of expression in iPSCs versus iPSC-CMs and visualized using ‘pheatmap’ (R). The average coverage per bin across gene categories was calculated, normalized to 0-1 rank, and then plotted using R (**Supplementary Fig. 3a**).

**Analysis of chromatin state annotations in ChIP-Seq and ATAC-Seq regions**. To compare the regulatory regions identified in our datasets with reference chromatin state maps, genomic coordinates identified by ChIP-Seq and ATAC-Seq experiments from iPSC or iPSC-CM meta-samples were intersected with the Roadmap Epigenomics 25-state ChromHMM annotations of 127 human tissues^40^ using ‘bedtools jaccard’. For each tissue and each chromatin state, the *Jaccard* similarity score measured the number of intersecting bases divided by the union of each pair of dataset. Enhancer chromatin states (‘9_TxReg’, ‘10_TxEnh5’, ‘11_TxEnh3’, ‘12_TxEnhW’, ‘13_EnhA1’, ‘14_EnhA2’, ‘15_EnhAF’, ‘16_EnhW1’, ‘17_EnhW2’, ‘18_EnhAc’) were selected. Similarity scores for each enhancer chromatin state were mean-centered normalized across all 127 tissues, ordered by mean similarity and hierarchically clustered and plotted using ‘pheatmap’ (**Fig. 1c-e and Supplementary Fig. 3b-c**).

**Normalization and analysis of variability of molecular phenotypes**. For RNA-Seq, we restricted the analysis to autosomal genes that had on average a minimum of 1 TPM per sample (14,933 and 15,167 genes analyzed respectively for iPSCs and iPSC-CMs) and integer-rounded RSEM expected counts were used as expression levels. For ChIP-Seq, we excluded peaks > 5 kb long and those located on sex chromosomes, resulting in 110,345 H3K27ac peaks analyzed in iPSCs, and 83,689 H3K27ac peaks and 37,994 NKX2-5 peaks analyzed in iPSC-CMs (**Supplementary Table 3**). Matrices of raw expression levels or peak coverage for each of the 5 datasets were vst-normalized using DESeq2 and analyzed for principal components (PCs) using R. To investigate the major sources of variability within each dataset, values for the first 10 PCs were correlated with known covariates across samples using ANOVA. For iPSCs, we tested association with sequencing batch, passage and subject, and for iPSC-CMs, we tested TNNT2 expression TPM, %TNNT2 positive cells, protocol of differentiation (day 15, day 15 + lactate purification, day 25 + lactate purification) and subjects. For iPSC-CMs, the sequencing batch coincided to the protocol of differentiation and was not included. For ChIP-Seq, we also included efficiency (measured as FRiP, fraction of reads mapping to peaks) as a covariate. The covariates that were most associated with PC1 were sequencing batch for iPSCs and TNNT2 expression and protocol for iPSC-CMs (**Supplementary Fig. 4**). We corrected the respective datasets by fitting a model including these covariates using the ‘lmFit’ function from ‘limma’ package and calculating the residuals using the ‘residuals’ function in R. Mean expression and coverage values for each gene/peak were then added back to the residuals. Residual-corrected values were then used in all subsequent analyses.

To assess the consistency of data generated from cell lines derived from the same individual versus cell lines from different individuals, we selected the 1,000 most variable genes or peaks and computed matrices of Spearman correlation values across all pairs of samples for each molecular phenotype. We then separated correlation values between pairs of samples from either the same or different individuals and calculated the average correlation per sample. Technical replicates were excluded for the comparisons between samples of the same subject. We tested for significant increase in correlation between samples from the same subject using a Mann-Whitney test (**Supplementary Fig. 4**). The same method was used to calculate correlation differences between samples from the same, related and unrelated subjects (**Fig. 2**).

**Allelic Specific Effect (ASE) analysis**. ASE analysis of RNA-Seq and ChIP-Seq data was performed as previously described^22^. To increase sensitivity of ASE and maximize the number of genes/peaks to analyze, reads from the two replicates of each individual per assay were merged. Heterozygous SNVs were identified by intersecting variant calls from WGS with either exonic regions from Gencode v19 or regions identified by each ChIP-Seq dataset. The WASP pipeline^88^ was employed to reduce reference allele bias at heterozygous sites. The number of read pairs supporting each allele was counted using the ASEReadCounter from GATK (3.4-46)^77^. Heterozygous SNVs were then filtered to keep SNVs where the reference or alternate allele had more than 8 supporting read pairs, the reference allele frequency was between 2-98%, the SNV was located in unique mappability regions according to wgEncodeCrgMapabilityAlign100mer track, and not located within 10 bp of another variant in a particular subject (heterozygous or homozygous alternative)^89–91^. ASE P-values for each SNV were calculated in each sample using a binomial test method described by^89,91^.

To combine ASE results at each SNV across samples, we performed a meta-analysis on all samples that were heterozygous for a given SNV and for which ASE could be tested. The binomial P-values of heterozygous SNVs were combined using the Stouffer z-score transformation method^92^. P-values were first transformed to z-scores with negative or positive sign according to the direction of the effect with respect to the alternate allele, then summed and transformed using the formula 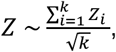 where Z is the z-score and *k* is the number of individuals for each SNV. The combined meta z-scores were transformed to P-values and a BH FDR was calculated using ‘p.adjust’ in R. The alternate allele frequency was averaged across all heterozygous samples. For downstream analyses, a cutoff or FDR<0.05 was used to define ASE SNVs.

**Correlation of ASEs across all individuals**. To evaluate if the allelic effect of ASE was consistent with the direction of homozygous effects within all family members, we performed linear regression on the gene expression or peak coverage as a function of genotype. Gene expression or peak coverage was normalized to *vst* and corrected for covariates using residuals as described above, and then normalized to z-scores across individuals. Genotypes were coded as 0, 1 or 2 for homozygous reference, heterozygous, and homozygous alternate respectively, and only one SNV per region (with the lowest combined ASE P-value) was used. A linear model in R ‘lm(z-score~genotype)’ was used to obtain the ß coefficient indicating the direction of effect. Spearman correlation was used to compare ß to the average allele proportion of the alternate allele to estimate the consistency of effects (**Fig 3d-f**).

**Correlation of ASEs across different molecular phenotypes**. To test if the direction of ASE of SNVs within ChIP-Seq peaks correlated with changes in peak coverage of other ChIP-Seq peaks or with gene expression, we performed a linear regression between the ASE-SNV genotypes and each phenotype. ChIP-Seq peaks were paired with the closest gene or peak within 500 bp using ‘bedtools closest’. Using linear regression, we tested the association between the individual genotypes (0, 1, 2) of the ASE-SNVs (FDR <0.05) and either the corresponding corrected and z-score normalized peak coverage or gene expression or those of the closest feature. In both peak/gene and peak/peak pairs, Spearman correlation coefficient was calculated between the two slopes (β) of linear regression. Only one ASE-SNV per peak/gene, peak/peak pair was considered. Spearman coefficient and BH corrected P-values were visualized using ‘heatmap.2’ (R) (**Fig. 3g-h**).

**Analysis of SNVs altering TFBS motifs**. The effect of NKX2-5 ASE-SNVs on TFBS motifs was estimated using position probability matrices (PPMs) of the 12 most enriched families of motifs identified using HOMER (**Supplementary Table 4**), from a library of known motifs. For NKX2, GATA, TEAD, MEF2, TBX20 and PDX1, we also used PPMs derived from a de-novo analysis. All PPMs are provided in **Supplementary Table 4**. Position weight matrices (PWMs) were calculated from the PPMs using a background nucleotide frequency of 0.25 for each base. To test whether SNVs with ASE altered a TF motif we used an approach similar to Ward et al.^44^. The sequences flanking each SNV that was tested for ASE (window of 40 bp) were obtained from the hg19 reference, and we created two sequences, one carrying the alternate and the other the reference allele. Using an in-house developed R script, we scanned each sequence with PWMs for each motif and identified the position with the highest score. For SNVs where either the reference or the alternate sequence matched or exceeded the log odds detection threshold reported by HOMER PPMs, the difference between the scores of the two alleles was calculated. In cases where a SNV matched multiple motifs from the same family, we kept only the motif with the highest score for either of the alleles. Fisher’s exact test was used to calculate enrichment for motif-altering SNVs in variants with ASE compared to variants without ASE. Enrichment was considered significant at BH corrected P-value <0.05 for multiple motifs testing (**Fig. 4a**). For ASE variants, the SNVs were tabulated according to whether the preferred ASE allele had a higher or lower motif score (**Fig. 4b**). For each of the 12 motifs we also calculated Spearman correlation between the allelic imbalance proportion of the reference allele and the difference in motif score between the reference and the alternate allele (**Fig. 4c-d and Supplementary Fig. 5**). For the most frequently disrupted motifs, the frequency by which each base of the motif was altered by an ASE-SNV was estimated (**Fig. 4e-h**). Motifs that were altered at NKX2-5 ASE-SNVs are indicated in **Supplementary Table 5**.

**Enrichment of ASE-SNVs for known quantitative trait loci**. To examine the enrichment of ASE-SNVs in known quantitative trait loci across different tissues, we obtained dsQTLs in LCLs from Degner et al.^43^; eQTLs from iPSCs from DeBoever et al.^22^, and eQTLs from HaploReg v4.1^44^, which contained combined results from 13 different including GTEx v.6^91^. All SNVs tested for ASE in ChIP-Seq datasets (H3K27ac in iPSCs and H3K27ac and NKX2-5 in iPSC-CMs) were intersected with these datasets and, for each annotation, we calculated the enrichment between heterozygous SNVs with and without ASE, using Fisher’s exact test in R. To identify tissue-specific eQTLs (**Fig. 5d**), the 44 tissues from GTEx were classified into 26 groups by merging similar tissues (adipose (n=2), artery (n=3), brain (n=10), cell lines (n=2), colon (n=2), esophagus (n=3), heart (n=2), skin (n=2), and the remaining 18 tissues were n=1). A gene-eQTL combination was defined as tissue-specific if 50% or more of the significant associations were in a single tissue group. Enrichment for eQTL annotations between significant and non-significant ASEs was determined by Fisher’s exact test BH corrected *P* value <0.05. In cases where multiple SNPs overlapped a peak, we counted only one SNP per peak.

**Enrichment of GWAS-SNPs in regulatory regions in iPSC-CMs**. To calculate enrichment for GWAS-SNPs in ChIP-Seq and ATAC-Seq peaks, we extracted sets of lead SNPs associated with 6 EKG traits from the GWAS catalog^93^: QT interval (matching ‘QT interval’ or ‘electrocardiography, QT interval’, 156 SNPs); PR interval/segment (matching ‘PR segment’, ‘PR interval’, ‘electrocardiography, PR interval’, or ‘electrocardiography, PR segment’, 52 SNPs); QRS duration (matching ‘QRS duration’, ‘QRS complex’, ‘QRS complex, QrS duration’, ‘QRS amplitude, R wave amplitude’, ‘QRS amplitude, QRS complex’, 80 SNPs); atrial fibrillation (matching ‘atrial fibrillation’, 73 SNPs); ‘heart rate (matching ‘heart rate’, ‘RR interval’ or ‘resting heart rate’, 175 SNPs); and P wave duration (matching ‘P wave duration’ and ‘electrocardiography, P wave duration’). To ensure that SNPs within each set were independent from each other, we pruned the lists of GWAS-SNPs using the ‘–clump’ function in PLINK v1.07^94^ with cut-offs of r^2^=0.5 and distance of kb=250. The SNP with lowest GWAS P-value was kept. We also extracted and pruned SNPs associated with 119 other traits that were associated with a similar number of SNPs (between 50 and 160 independent SNPs per trait), which are listed in **Supplementary Fig. 6**. We used GREGOR^95^ to test each of these 125 SNP sets for enrichment in ChIP-Seq and ATAC-Seq peaks from iPSCs and iPSC-CMs from this study as well as in peaks from cardiac tissues from Roadmap (fetal heart DHS, right ventricle and right atrium H3K27ac), used as positive controls. An r^2^> 0.8 (1000 Genomes phase 1 EUR) was used to include LD proxies and 500 random null sets matched on minor allele frequency and number of SNPs were used to calculate P-value and fold-enrichment over expectation based on a binomial distribution^95^. Enrichment P-values were BH FDR corrected for multiple testing using ‘p.adjust’ (R).

To calculate the enrichment for EKG GWAS-SNPs in NKX2-5 ASE-SNVs, we used GREGOR to obtain SNVs overlapping NKX2-5 peaks and associated with any of the six EKG traits (417 independent lead variants plus variants in LD r^2^>0.8 in EUR). For the SNVs that could be tested for ASE (heterozygous and a minimum of 8X read depth), we calculated the proportion with and without ASE and tested their relative enrichment using the Fisher exact test.

In **Supplementary Table 5**, we also show overlap between all GWAS-SNPs in the NHGRI-EBI catalog (all traits) and 3,117 ASE-SNVs found in ChIP-Seq of NKX2-5 (1,941), H3K27ac in iPSC-CMs (803) and H3K27ac in iPSCs (373). To annotate these variants we obtained all lead SNPs from the NHGRI-EBI GWAS catalog accessed on 10/20/2017^93^ and extracted SNPs in LD with them (r^2^>0.8) in Asian (ASN) and European (EUR) populations from the 1000 Genomes Project Phase 1^96^.

**Estimating heart rate GWAS enrichment in molecular phenotypes and prioritizing putative causal variants**. We employed the fgwas framework^55^ to integrate the iPSC-CMs molecular phenotypes and EKG GWAS data in order to identify putative causal variants and novel associations. We obtained summary statistics from the den Hoed et al.^6^ heart-rate GWAS meta-analysis (2,516,407 SNPs analyzed) from LD hub (http://ldsc.broadinstitute.org/ldhub/). To determine the enrichment of genetic variants influencing heart rate within the different iPSC-CM molecular phenotypes, we first annotated each variant with the type of molecular phenotype it overlapped: peaks (ATAC-seq, H3K27ac, and NKX2-5 peaks) and/or ASE-SNVs (H3K27ac and NKX2-5). We then applied fgwas on these 5 annotations, using the *β* and standard error of the *β* from the GWAS as inputs, to consecutive ~1Mb intervals across the genome (-k 900). We followed the recommended fgwas pipeline: briefly, we first calculated enrichment for each annotation individually, and then added annotations into the model one at a time (jointly modeling them) keeping the annotation that increased the likelihood of the model the most, until no more annotations remained or increased the likelihood. Next, we determined the best cross-validation penalty by trying penalties between 0.01 and 0.30 in steps of 0.01, settling on a penalty of 0.11 as it had the highest likelihood. The final estimated ridge parameters, reported as the ln(Enrichment), were: NKX25-ASE-SNVs: 2.762, H3K27ac peaks: 1.549, NKX2-5 peaks: 1.03, ATAC: 0.465, H3K27ac ASE-SNVs: −0.256. As the model with all 5 annotations had a better likelihood than any model with 4 annotations, we used the full model with fgwas to update the Bayes Factors for each variant using the cross-validation estimated ridge parameters, and calculated the posterior probability of association (PPA) for each variant within the same consecutive 1Mb windows used above. The PPA is the proportion of the total GWAS risk signal at a locus measured by Bayes Factors that is attributed to a particular variant, multiplied by the probability that the genomic region contained a real signal (either estimated by fgwas for novel loci, or set to 1 for known loci). We selected variants with PPA>0.3 (21 variants, **Supplementary Table 6**).

**Hi-C**. We used the Hi-C data from from 13 iPSC-CMs samples generated in this study (**Supplementary Table 2**) and described in detail in Greenwald et al^58^. Briefly, *in situ* Hi-C was performed on 2-5 million cells as previously described^97^. Formaldehyde-crosslinked cells were lysed and nuclei were digested with 100U MboI overnight at 37°C. Fragmented ends were biotinylated for 90 min at 37°C, diluted and proximity ligated for 4 hours at room temperature. Crosslinks were reversed and samples were then purified by ethanol precipitation, resuspended in 100uL 1X Elution Buffer, fragmented using a Covaris S2 instrument, and size selected using AmpureXP beads. Subsequently, biotinylated ligation junctions were pulled down using T1 Streptavidin beads and used for library preparation consisting of end-repair, dA-tailing, adapter ligation, PCR amplification and purification. Libraries were sequenced on an Illumina HiSeq 4000 using 150bp paired-end reads.

For each sample, Hi-C reads were aligned to human reference genome hg19 using BWA-MEM (version 0.7.15) with default parameters, with forward and reverse reads from the paired-end data being aligned independently. Paired-end reads were then reconstructed, processed, and filtered using the Juicer pipeline^98^. These filtered read pairs (contacts) were pooled across all iPSC-CM samples to obtain a higher resolution chromatin contact map. Loops were called with a combination of two distinct loop calling algorithms, HICCUPS^97^ and Fit-Hi-C^99^, and subsequently filtered to high confidence loops with pgltools^100^, resulting in 19,003 high-resolution chromatin loops with a median size of 403 kb.

**Gene expression analysis of 128 iPSC-CMs**. To determine if candidate functional NKX2-5 ASE-SNVs overlapping EKG traits GWAS were associated with gene expression, we used RNA-Seq data of iPSC-CMs differentiated from iPSCs of 128 different individuals of the iPSCORE cohort^36^, which are part of an ongoing study. Subjects included 43 males and 85 females, of age between 9 and 88, of diverse ethnicities (with the majority being Europeans, n=78, and Asians, n=23) and all have 50X WGS from blood. iPSCs were differentiated into day-25 cardiomyocytes using the method described above, including a 4mM Sodium L-Lactate enrichment step at day 15, and yielded on average 83.9 +/− 13.6 % cTNT positive populations. RNA-Seq was generated and processed using the same pipeline as described above. Raw gene expression data were first filtered for genes with TPM ≥ 2 in at least 5% of the samples and then quantile-normalized. From these values we then calculated PEER factors^101^ and used the residuals of the first 10 PEER factors as normalized gene expression values. We then extracted the individuals’ genotypes and the normalized gene expression values and performed linear regression for the specific SNP-gene associations in R (**Fig. 7**).

## Data Availability

All iPSC lines are available through WiCell Research Institute (www.wicell.org; NHLBI Next Gen Collection). Phenotype and array genotype data is available through dbGaP (phs000924). Sequence data analyzed in this study will be deposited in the NCBI Sequence Read Archive (SRA, http://www.ncbi.nlm.nih.gov/sra) and accessible through dbGaP (phs000924).

## Code Availability

Computational analyses have been performed using the indicated publicly available software, as well as R and python scripts developed by the authors. All codes are available upon request.

## Acknowledgements

This work was supported in part by a California Institute for Regenerative Medicine (CIRM) grant GC1R-06673, and NIH grants HG008118-01 and HL107442-05. PB is supported by the Swiss National Science Foundation Postdoc Mobility fellowships P2LAP3-155105 and P300PA-167612; CD was supported in part by the University of California, San Diego, Genetics Training Program through an institutional training grant from the National Institute of General Medical Sciences (T32GM008666) and the California Institute for Regenerative Medicine (CIRM) Interdisciplinary Stem Cell Training Program at UCSD II (TG2-01154). Library preparation and sequencing services were conducted by Kristen Jepsen and Mahdieh Khosroheidari at the UCSD IGM Genomics Center supported by NIH grant P30CA023100. We are very thankful to Chia-An Yen and Nathanael Spann for assistance with ChIP-Seq experiments and to Anthony Schmitt for Hi-C data. We thank many colleagues for helpful comments.

## Author contributions

PB generated the experimental data and performed statistical analyses. ADC generated iPSC-CMs and performed data generation. WWG processed and analyzed Hi-C data and performed the fgwas analysis. CD performed RNA-Seq and ATAC-Seq processing and implemented the allele-specific effect pipeline. HL processed WGS, ChIP-Seq and Hi-C data. FD and SS generated iPSC-CMs and participated in data generation. HM assisted with data processing and computational analyses. MD and ENS participated in statistical analyses. KAF conceived and oversaw the study. PB, KAF and ENS prepared the manuscript.

